# A single evidence accumulation process informs perceptual choices and subsequent confidence reports

**DOI:** 10.1101/2025.06.05.658071

**Authors:** J.P. Grogan, L. Vermeylen, S.L. Mannion, C. McCabe, D. Monakhovych, K. Desender, R.G. O’Connell

**Author notes:** the authors contributed equally to the work (J.P. Grogan, L. Vermeylen). the authors jointly supervised the work (K. Desender, R.G. O’Connell).

## Abstract

How is the content of our decision making processes translated into subjective confidence? Current computational models designed to account for choice- and confidence-reporting behaviour disagree on two fundamental questions: do choice and confidence reports arise from shared versus distinct evaluation processes and what stopping-rule is applied to evidence sampling for confidence reports? Here, we employed a neurally-informed modelling approach in which competing models were compared both based on their fits to behaviour and their ability to recapitulate human electrophysiological signatures of evidence accumulation. Participants made self-paced confidence reports after indicating perceptual choices on a random dot motion task. Model comparison based on behavioural data alone showed that boundary-based stopping rules for confidence reporting provided superior fits than time-based stopping rules. However, results were inconclusive when comparing models with a single evidence accumulation process dictating choice and confidence reports compared to models invoking distinct processes. Importantly, the single-process model markedly outperformed the other models in its ability to account for the observed neural evidence accumulation patterns, including the fact that signal amplitudes predicted reported confidence several hundred milliseconds before the initial choice had been reported. Our study demonstrates that choice behaviour, subjective confidence, and associated neural decision signals can be jointly explained by a model invoking a single process of evidence accumulation with separate boundaries for the initial choice and subsequent confidence reports.

**Significance Statement:** How does the brain generate representations of subjective confidence from categorical decisions? While some models suggest that confidence arises from a read-out of the same information that determined the initial choice, others propose that confidence is determined by a distinct computational process. Definitively testing these alternative accounts has proven difficult because they often make similar predictions for behavioural data. Here, we employ a novel neurally-informed modelling method that evaluates models based on their ability to reproduce key neurophysiological signatures of choice and confidence in addition to behaviour. Our results indicate that choices and subjective confidence are determined by a shared sensory evidence accumulation process.

## Introduction

Convergent evidence from neurophysiological recordings and mathematical models suggests that the evidence accumulation processes that underpin decision formation also contribute to our subjective sense of choice confidence. How and when those processes contribute to confidence remains unknown. In most human studies of choice confidence, participants are asked to make an initial categorical decision before reporting their confidence in that choice. These metacognitive reports tend to exhibit a number of systematic features including decreased confidence, slower confidence response times and increased change-of-mind probability following incorrect decisions (1–3). A variety of different computational models have been devised to account for these behavioural patterns, most of which invoke some form of evidence accumulation that gives rise to the reporting of confidence (4–9) While neurophysiological data do point to the operation of evidence accumulation processes prior to metacognitive reports, both in the period leading up to a choice (1, 10–13) and in the interval between choice and metacognitive reports (12, 14–16), there is ongoing uncertainty regarding several fundamental elements of this process.

The first key element that existing models disagree on is whether or not choices and confidence arise from a shared neural currency. Under one view, the same evidence accumulation process that drives the initial choice also drives confidence judgments (4, 10, 13), with some models allowing for evidence accumulation to continue after choice commitment to inform delayed confidence reports (4). This view is supported by representations of confidence emerging in decision-related evidence accumulation signals before (11–13, 17) and after commitment (12). An alternative view is that choices and confidence are driven by distinct meta-cognitive evidence accumulation processes (14, 18). For example, some studies have suggested that evidence accumulation processes giving rise to confidence operate within a distinct reference frame, selectively accumulating evidence that an error has been made (14, 16), or that confidence is informed by an additional parallel accumulation process integrating information about stimulus reliability (8). Going one step further, recent work argued that the output of the confidence accumulator is used to online control the height of the decision boundary during decision formation (9). In sum, whether evidence accumulation processes giving rise to confidence are qualitatively distinct from those giving rise to confidence remains unclear.

A second element that computational models disagree on, is the stopping-rule that is applied to the accumulation process giving rise to confidence. Regardless of whether the accumulation process for confidence is shared with that for choices or reflects a qualitatively different process, models require a stopping rule determining when to quantify confidence. One class of model assumes that evidence is accumulated to inform confidence for a fixed duration on each trial (with that duration varying across trials) after which the total accumulated evidence is compared against confidence criteria to generate explicit confidence ratings (4, 8). Alternative models propose that accumulation for confidence only ceases when the decision variable reaches an additional set of confidence boundaries that come into play once initial-choice commitment has been reached. These confidence boundaries usually collapse over time, ensuring that later confidence reports will be associated with lower confidence(6, 7). In addition, some confidence models do not specify any stopping rule and therefore cannot account for systematic variations in the timing of self-paced reports(9, 19).

The extent to which these models can be distinguished based solely on fits to behavioural data likely depends at least in part on whether and how the initial choice data are modelled. In keeping with the dominant models of perceptual choice (e.g. Drift Diffusion Model), most models of post-choice confidence assume that the initial choice is subject to a constant bound (4–7). This has the consequence that the same amount of evidence will have been accumulated by the time of commitment for all choices. As a result, models invoking a distinct accumulation process for confidence versus a single continued accumulation process tend to make similar predictions for confidence reports. However, there is now strong evidence from computational modelling and neurophysiology that observers collapse their choice boundaries as time elapses when choices must be reported under time pressure (20–24). One consequence of having collapsing choice boundaries is that cumulative evidence at choice commitment can vary substantially and systematically across trials, which could provide a crucial additional constraint on the nature of the evidence accumulation signal underlying confidence.

Our goal in the present paper was therefore to jointly model perceptual choices and subsequent self-paced confidence reports, and to compare models with different stopping-rules and different accumulation processes underlying confidence. In addition to evaluating our models based on their fits to behaviour, we examined the extent to which they could recapitulate the observed pre- and post-choice dynamics of a well-validated neural signature of evidence accumulation known as the centro-parietal positivity (CPP). The CPP is a component of the human event-related potential that exhibits a gradual, evidence-dependent, RT-predictive build-up during decision formation, irrespective of whether the choice entails an immediate response, a delayed response, or no overt response at all (24–27). The CPP’s pre-choice amplitude is modulated by contextual factors such as prior knowledge and time pressure, consistent with boundary adjustments identified by behavioural models(24, 28, 29). It declines as a function of time alongside choice accuracy, consistent with a collapsing bound effect(24, 29), and scales with explicitly reported confidence (12, 17, 30–33). Recent work has also established that the CPP continues to evolve after an initial choice report and that its post-choice amplitude predicts subsequent cued confidence reports (12). To our knowledge, no previous study has examined the CPP’s dynamics prior to self-paced confidence reports which may be essential in order to identify the stopping rules and evidence accumulation processes being implemented.

## Results

### Post-decision deadlines induce a Confidence-SAT

We recorded 128-channel EEG during a two-alternative random-dot motion discrimination task (see Figure 1). Participants reported the direction of coherent motion within a 1500ms deadline after which the stimulus remained on screen until participants rated their confidence on a 6-point scale (‘certain left’ to ‘certain right’). In order to facilitate identification of the stopping rule for post-choice accumulation, we manipulated the time pressure applied to the post-choice confidence reports by varying the deadline for response (Speed Regime = 700ms; Accuracy Regime = 3000ms). We defined ‘confidence’ as the degree to which the confidence report aligned with the initially chosen alternative (e.g., after a left choice, the highest confidence level would be ‘certain left’ and the lowest confidence level would be ‘certain right’) and ‘certainty’ as the participant’s confidence irrespective of whether or not they were endorsing the initial choice (e.g., after a left choice the highest certainty levels would be ‘certain left’ and ‘certain right’). Response times (RT) for the confidence reports (confidence RT) were measured relative to initial choice RT. We also measured ‘final-accuracy’, the extent to which the confidence report indicated the correct motion direction; ‘metacognitive sensitivity’, the degree to which confidence ratings distinguished correct from incorrect responses (AUROC2, see Methods), and ‘changes-of-mind’, defined as trials on which the confidence report indicated that the incorrect alternative had initially been chosen.

**Figure 1.**
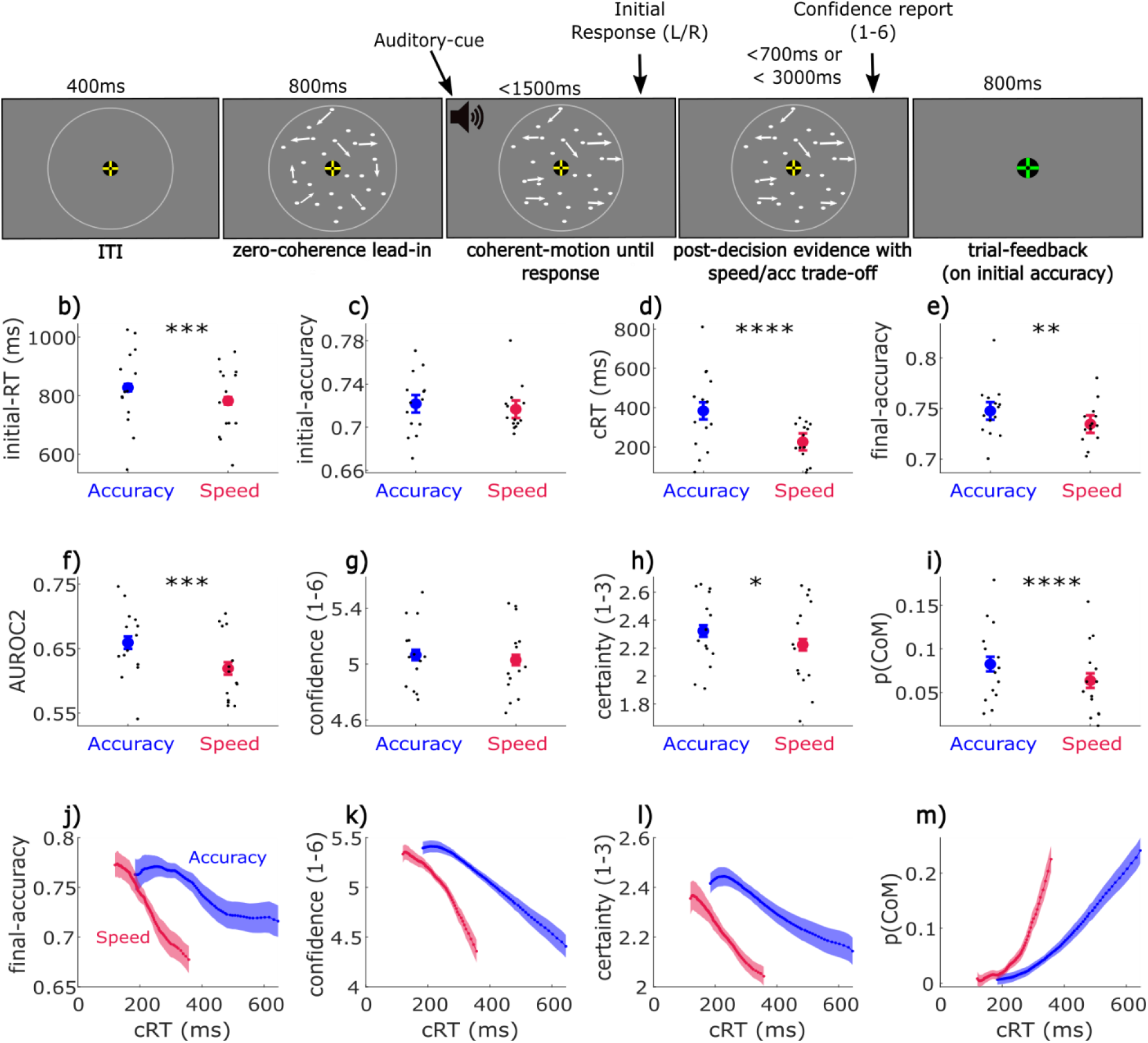
Task design and Speed-pressure effects on behaviour. a) Random-dot motion task with zero-coherence lead-in, before a 1500ms deadline for an initial left/right choice. Moving dots with the same coherence remain on the screen as post-decision evidence, for either 700ms or 3000ms (blocked design) to induce a post-decision speed-accuracy trade-off; participants must rate their confidence before this deadline. b-k) Behavioural results; blue/red dots show overall means with 95% confidence intervals, black dots show individual-participants’ means, and black stars show the significance of the speed-pressure effect, see text for full statistics (*=p<.05, **=p<.01, ***=p<.001, ****=p<.0001). Speed-pressure significantly decreases initial RT (b), confidence-RT (cRT; d), final-accuracy (e), AUROC2 (f), mean certainty (h) and changes-of-mind (CoM) (i). It did not significantly decrease initial-accuracy (c) or mean confidence (f). j-m) Conditional plots showing sliding window analysis of speed-pressure effects. Within each person and condition, we took a sliding window of 20% of cRTs, from the 1^st^ to the 80^th^ percentile, and plot the mean y-axis value at the mean cRT for that percentile-window, across participants, with shading showing SEM. This measures the change in the y-axis measure over increasing percentiles of cRT, while accounting for individual differences in mean and range of cRTs. Slower cRTs were associated with lower final-accuracy (j), lower confidence (k), lower certainty (l) and more changes-of-mind (m), with steeper slopes in the Speed-pressure condition.

Despite the fact that the physical evidence remained available during the interval between the choice report and the confidence-reporting deadline, the average confidence-RTs were overall quite fast in both Regimes (Speed: Mean = 226ms, SD = 96ms; Accuracy: Mean = 384ms, SD = 204ms). As expected, initially correct choices had faster initial-RT (log RT: β=-0.1091, t(1,25331)=-8.2343, p<.0001), faster confidence-RT (log confidence-RT: β=-0.0905, t(1,25331)=-4.2684, p<.0001), higher certainty ratings (β=0.1483, t(1,25331)=7.0579, p<.0001), higher confidence ratings (β=0.2607, t(1,25192)=8.4610, p<.0001), as well as fewer changes-of-mind (logistic regression: β=-0.7821, t(1,25323)=-33.6550, p<.0001). Participants reported their confidence more quickly in the Speed Regime (Figure 1d; single-trial mixed-effects regression on log(confidence-RT): β=-0.3494, t(1,25331)=-4.9135, p<.0001) and with lower final-accuracy (Figure 1e; logistic regression: β=-0.0408, t(1,25331)=-2.8443, p=.0045). Overall confidence ratings were not significantly affected by the speed pressure manipulation (Figure 1g; β=-0.0162, t(1,25224)=-1.0173, p=.3090) but there was a significant reduction in certainty (Figure 1h; β=0.0637, t(1,25331)=-2.4627, p=.0138). Although the speed-pressure manipulation was only applied to the post-choice confidence reports, this nevertheless resulted in a slight decrease in initial RT (Figure 1b) and accuracy (Figure 1c), although only the effect on RT reached statistical significance (single-trial regression of log(RT): β=-0.1196, t(1,25331)=-3.7660, p=.0002; logistic regression on accuracy, β=-0.0187, t(1,25331)=-1.3342, p=.1822). In addition, there was a significant reduction in the frequency of changes-of-mind in the Speed Regime (Figure 1i; logistic regression: β=-0.1481, t(1,25331)=-6.0496, p<.0001); as changes-of-mind are usually corrective, this explains the lower final-accuracy in the Speed regime, as initial-errors were not corrected as often. Despite there being no overall reduction in confidence, metacognitive sensitivity was significantly reduced in the Speed Regime (Figure 1f; aggregate level regression: β=-0.3630, t(1,26)=-4.2433, p=.0002), replicating the findings of Herregods et al. (6).

Additionally, we observed that longer confidence-RTs were associated with lower final-accuracy (β=-.1465, t(1,24329)=-9.272, p<.0001), lower confidence (β=-0.3887, t(1,25222)=-5.2321, p<.0001) and certainty ratings (β=-0.1663, t(1,25239)=-4.0614, p<.0001), and more changes-of-mind (β=1.0922, t(1,25329)=12.9930, p<.0001), and these relationships were stronger in the Speed Regime (speed-pressure*confidence-RT on final-accuracy: β=-0.0560, t(1,25239)=-3.5577, p=.0004; confidence: β=-0.0967, t(1,25222)=-3.0604, p=.0022; certainty: β=-0.0549, t(1,25329)=-2.0666, p=0.0388; changes-of-mind: β=0.2739, t(1,25329)=9.8300, p<.0001). This pattern is consistent with confidence reports being subject to collapsing boundaries.

### Comparing Model fits to Choice and Confidence Reports

To better understand the cognitive and neural mechanisms underlying perceptual choices and subsequent confidence judgements we turned to evidence accumulation models. To this end, we used a two-stage fitting procedure where we first fit a variant of the DDM in which the decision boundaries were free to linearly decrease over time, and then incorporated those best-fitting parameters into models that were designed to also account for the final confidence reports (note that a two-stage fitting approach was adopted because it was found to significantly improve parameter recovery compared to fitting all parameters at once). The initial choice data were well explained by the model (Figure 2a, Extended Data Figure 1) which identified a substantial degree of boundary collapse that was common to both confidence report Regimes as well as a small reduction in initial boundary height in the Speed Regime (Extended Data Table 1).

**Figure 2.**
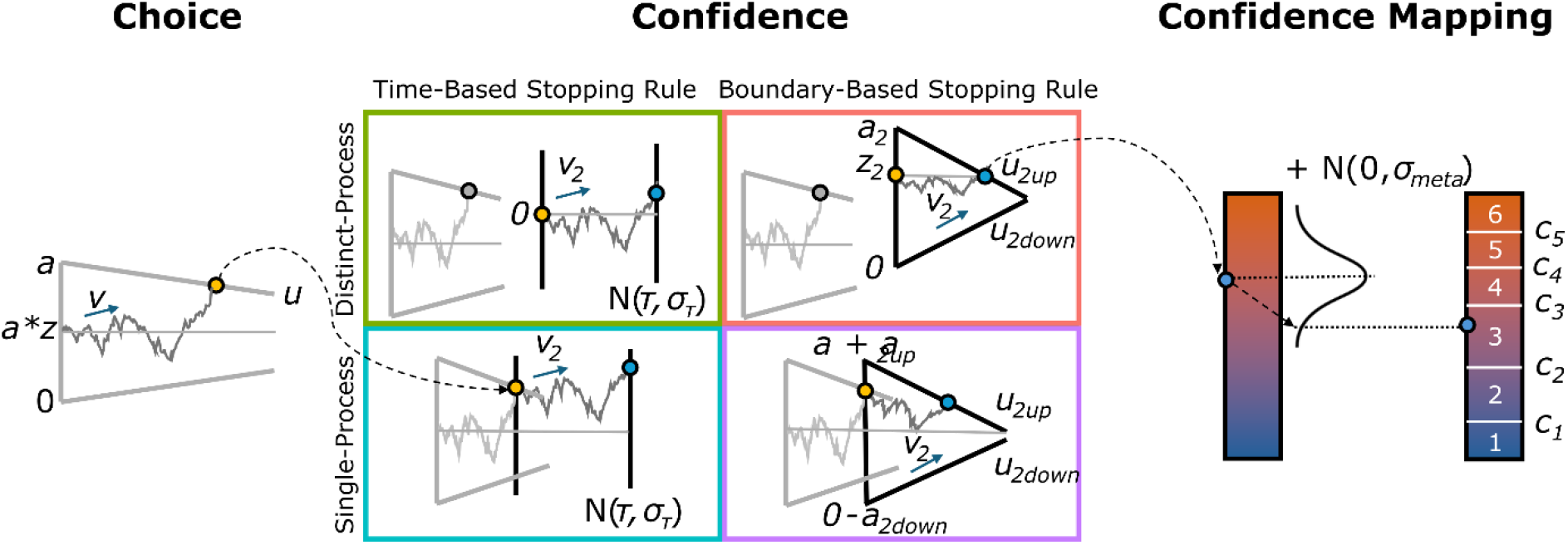
Overview of the Candidate Models. (left) The initial choice was modelled using a drift diffusion model, where noisy evidence accumulates with a certain drift rate (v) until hitting a collapsing boundary (a; collapse rate controlled by u). Accumulation started at the unbiased point a*z. (middle) In Time-based models, the time of stopping was determined by a mean deadline time (τ) with some variability (σ_τ_). In boundary-based models, the time of stopping was determined by collapsing boundaries, controlled by parameters related to confidence boundary heights (a_2up_ and a_2down_ for the Single-Process variant, a_2_ and z_2_ for the Distinct-Process variant), and corresponding collapse rates (u_2up_, u_2down_). Single-Process models evaluate evidence relative to the initial reference frame used during choice formation (i.e., the unbiased starting point), while Distinct-Process models evaluate that evidence from a new reference frame (i.e., the freely estimated confidence starting point z_2_). All models included a parameter for the rate of accumulation for confidence (v_2_). (right) Finally, in all models, evidence was translated into a six-point confidence scale using a metacognitive noise parameter (σ_meta_) and five confidence criteria (c1 to c5). Non-decision components not displayed (see Methods).

To investigate the mechanisms underlying the computation of confidence, we evaluated four classes of models (Figure 2, see Methods for details). In all models, the evidence used to compute confidence is accumulated by a decision variable (DV) whose level at the time of commitment to the confidence report is translated into a confidence rating by comparison against confidence-criteria. The models included free parameters to estimate the position of each of the confidence criteria, as well as a metacognitive noise parameter to capture trial-to-trial fluctuations in the mapping of the DV to these criteria. All models had separate choice- and confidence drift rates and non-decision time parameters (the latter was necessary in Single process models to account for additional motor delays associated with the confidence reports). Note that for the accumulation process governing confidence the non-decision time parameters were free to take on negative values, making it possible for the Distinct models to initiate confidence-related accumulation while the initial choice process was still ongoing. These non-decision times were estimated separately for changes-of-mind and no-change responses, to capture motor execution costs of switching hands between choice and confidence responses for change-of-mind trials.

The four models were distinguished based on two key categories: Time or Boundary based stopping rules; and Single or Distinct processes for choices and confidence (Figure 2). Time-based stopping rules allow the accumulation process for confidence to continue for a set amount of time on each trial, with that time being drawn from a gaussian distribution, while we followed the approach of Herregods et al., (2025) by allowing Boundary-based models to have linearly collapsing upper and lower confidence boundaries that terminate the process when reached. Having independent collapse rates for the upper and lower confidence boundaries allow these models to capture the full range of association between confidence and cRTs seen in empirical data (6). In Single-process models the accumulation process for confidence takes the form of a continuation of the pre-choice process, and therefore the DV’s proximity to the confidence boundaries at the time of initial choice commitment is determined by how much evidence has been accumulated up to that point. In Distinct-process models, the starting-point (and therefore proximity to the confidence boundaries) of the accumulation process for confidence is freely fit independent of the DV’s level at initial choice commitment, and the accumulation for confidence therefore only represents evidence accumulated *independent from* the initial choice.

We estimated the parameters of the four models by minimizing a composite objective function that included the distribution of confidence RTs conditional on decision accuracy, the proportion of confidence ratings conditional on decision accuracy, and the proportion of confidence ratings conditional on confidence-RT quantiles (see Methods for details). Parameter and model recovery analyses showed that the four models are dissociable from each other using BIC model comparison, and that parameters of individual models are recoverable (see Methods/Model Fitting Procedure & Extended Data Figure 6). Each model was fit separately to data from both the Speed and Accuracy conditions. The two-stage fitting approach ensured that the models were able to capture the relevant behavioural patterns for both choice and confidence, while ensuring that the four models only differed in terms of their accumulation processes for confidence.

Model comparison based on goodness-of-fit statistics accounting for model complexity indicated that Boundary-based models clearly outperformed the Time-based models (mean ΔBIC = 28.5; see Table 1). In particular, the collapsing Boundary models provided excellent fits for the shape of the confidence RT distributions (Figure 3A) and accurately captured that confidence RTs are much shorter than their choice RT counterparts (Figure 3C). The collapsing confidence bounds ensured that slower confidence RTs were associated with less accumulated evidence and therefore lower certainty, whereas this association was completely missed by the Time-based models (Figure 3B). The two Boundary models could not be conclusively distinguished in terms of their fits to behaviour, with only a slight preference for the Distinct over Single process model (ΔBIC = 3; Figure 3B&C). Both models captured the speed pressure effects on confidence via faster collapse of the upper confidence boundary (Boundary-Distinct: t(13) = -4.21, p = 0.0010; Boundary-Single: t(13) = -5.16, p = 0.0002) but disagreed regarding the relative magnitude of choice drift rate versus the confidence drift rate (with Boundary-Single indicating larger confidence drift rates and Boundary-Double indicating larger choice drift rates) and the role of non-decision time adjustments (with only Boundary-Single indicating a shortening of confidence non-decision time for non-change-of-mind trials in the Speed Regime; see Extended Data Table 2-3 for full parameter results).

**Table 1.**
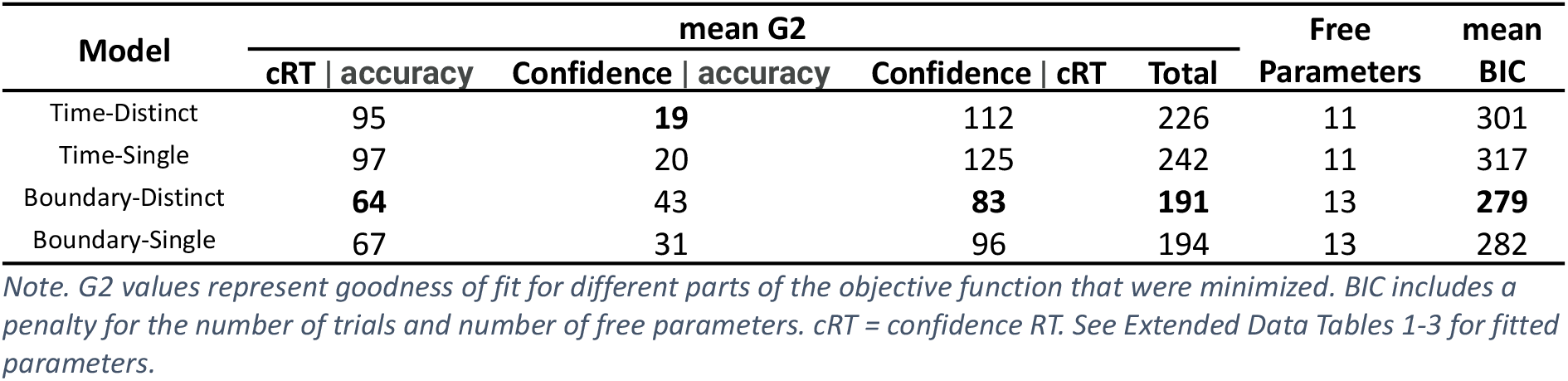
Goodness-of-fit metrics for the four classes of models. Values are mean scores over 14 participants in 2 conditions.

**Figure 3.**
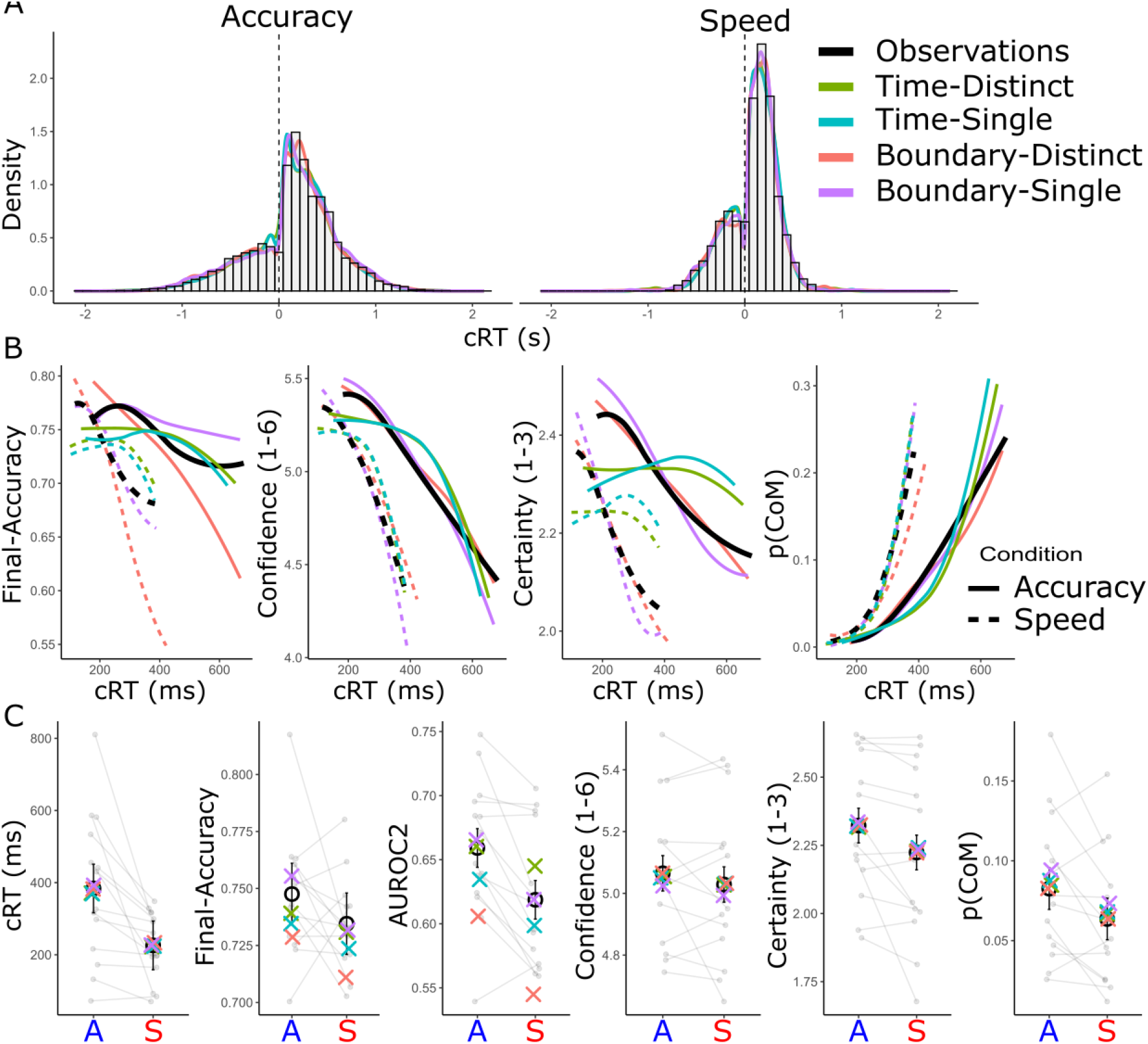
Model Fits to Behavioural Data Across Accuracy and Speed Conditions. (A) Distributions of confidence response times (cRT) separate for correct (positive RTs) and incorrect (negative RTs) initial-choices in the Accuracy (left) and Speed (right) conditions. (B) final-accuracy, confidence, certainty, and change-of-mind (CoM) probability as a function of confidence-RT (cRT; smoothed using loess regression). (C) Mean cRT, final-accuracy, metacognitive sensitivity (AUROC2), confidence, certainty, and CoM probability for the Accuracy (A) and Speed (S) conditions. Error bars denote the within-subject 95% CI. Black histograms, lines, and points represent observed data, while coloured densities, lines and crosses represent the various model predictions. CoM=Change-of-mind.

Although both Boundary models were nearly indistinguishable in terms of their overall fits, there were some qualitative differences in their predictions for behaviour. The Distinct-process variant benefits from greater flexibility due to the post-choice starting point being freely estimated, rather than being constrained by the DV’s level at initial choice commitment. This allows the Boundary-Distinct model greater freedom to adjust its post-decisional dynamics to match the observed confidence patterns. However, this greater independence from the initial choice process comes at the cost of lower than observed metacognitive sensitivity (AUROC2) and final-accuracy (Figure 3C). In contrast, the starting-point and confidence boundaries of the Boundary-Single model are directly determined by the termination point of the initial decision, providing a better fit to final-accuracy and metacognitive sensitivity (Figure 3C) and the relationship between confidence-RT and final-accuracy (Figure 3B) but at the cost of a slightly worse replication of the confidence-RT distributions.

In order to further distinguish between the models, we leveraged our concurrently recorded EEG data to compare their ability to recapitulate the pre- and post-choice dynamics of a neural proxy of evidence accumulation; the Centro-Parietal Positivity (CPP).

### Pre- and Post-choice neural evidence accumulation dynamics

In keeping with previous observations, the CPP showed a ramp-to-peak pattern leading up to the initial choice (Figure 4 - top row). The CPP is positive-going irrespective of which alternative the sensory evidence favours and, correspondingly, our previous work has shown that its amplitude at initial choice-commitment scales with the certainty of simultaneous confidence reports, irrespective of whether or not the confidence report indicates that an error had been made (i.e., Certainty rating; (12)). Here we found that the same relationship holds when confidence is reported *after* the choice, with the CPP reaching significantly higher amplitudes in the lead up to the initial choice and immediately prior to the final confidence report on trials rated with higher certainty (Figure 4A; black bars show significant clusters, p<.05). Note that confidence reports in our study were rather fast on average (Figure 3C), such that the choice-locked and confidence-locked CPP epochs overlap substantially, leading them to exhibit similar waveforms that are slightly shifted in time. We also found that speed-pressure affected the CPP but in the opposite direction expected from previous reports (24, 29), reaching a higher amplitude before the initial and confidence responses in the Speed Regime (Figure 4C - top row; black bars show significant clusters, p<.05). Next, we compared the two Boundary models’ ability to replicate these CPP effects.

**Figure 4.**
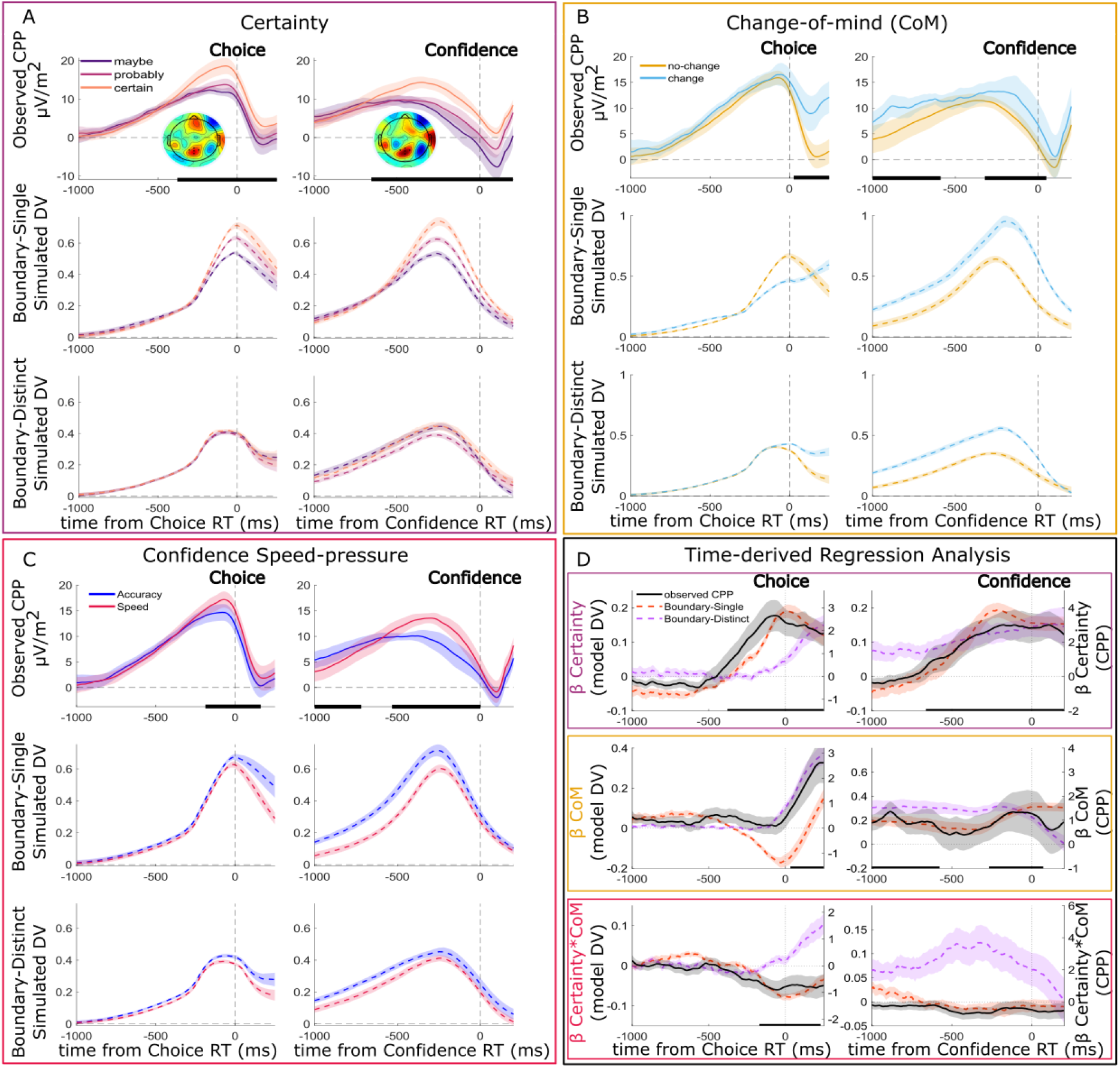
Observed CPP & Simulated Decision Variable, and Regression Analysis. (A) Certainty Ratings. (B) Change of mind (CoM). (C) Speed Pressure. (D) Time-derived regression analyses. In panels A-C, the CPP is presented in the top rows, the simulated decision variable from the Boundary-Single model in the middle rows and the Boundary-Distinct model in the bottom rows. Left columns show choice-locked waveforms and right columns show confidence-locked waveforms. Choice-locked epochs end 250ms after initial response, which is just longer than the average median confidence-RT (243ms). Topography insets show the effects of each factor on mean observed EEG activity -150:-50ms before the initial choice or the confidence response (for Certainty, the difference is between Maybe and Certain ratings);lack dots show the CPP electrodes. (D) Time-derived Regression analyses, showing the beta-coefficients for Certainty, CoM and Certainty*CoM effects for the model DVs (left-axis) and CPP (right-axis). Solid black bars along the bottom show clusters of significant effects on the observed CPP (single-trial time-derived LME regression, with permutation testing and cluster-control of FWER at .05) and Error bars reflect the within-subject SEM.

### Comparing Model and Neural Evidence Accumulation Time-courses

We used the best-fitting parameter values of each model to simulate evidence accumulation time courses. In keeping with previous work, since the CPP is positive-going irrespective of which alternative the evidence favours, we simulated the absolute value of accumulated differential evidence (24). As detailed in Methods, the simulations for all four models incorporated several additional features to match known properties of brain signals like the CPP. These features included modelling a noisy baseline period before evidence onset, separating non-decision time into stimulus encoding and motor execution components, and incorporating a post-commitment signal decay (see Methods and Extended Data Figure 2). The simulation of the post-choice CPP differed for the Single- and Distinct-process models, reflecting their contrasting hypotheses regarding the reference frame for evidence accumulation. In the Single-process models, evidence accumulation continued seamlessly from the initial decision stage. In Distinct-process models, however, a separate post-decisional accumulation process was initiated; this new process used a novel starting point, which served as a new reflection point for calculating the absolute value of differential evidence.

To distinguish between the behaviourally similar Boundary-Single and Boundary-Distinct models, we compared the simulated DV to the observed CPP in two ways (simulations for the Time-based models can be found in Extended Data Figure 3). First, we examined average waveforms separately for different certainty ratings, change-of-mind occurrences, and speed Regimes; and second, we used time-based regression analyses to more formally disentangle the effects of certainty and changes-of-mind. Both models captured the qualitative patterns observed in the grand-average CPP, including a rise-to-peak before the initial decision, similar choice- and confidence-locked waveforms, and a decay to baseline that starts several hundred milliseconds before the confidence report (Figure 4). Both models did a good job of matching the average CPP dynamics but the Boundary-Single model was markedly better at reproducing the observed amplitude effects of Certainty including their emergence several hundred milliseconds before the initial choice (Figure 4A) while the Boundary-Distinct model predicted very small Certainty effects. The two models performed similarly well when accounting for changes-of-mind in the post-choice interval, although the Boundary-Distinct model provided a slightly superior match to the amplitude differences observed around the time of the initial choice (Figure 4B).

We also examined the extent to which the models could reproduce the effects of post-choice speed pressure on CPP amplitude but found that neither model agreed with the empirical data, with both predicting larger amplitudes for the Accuracy compared to the Speed condition (Figure 4C).

As certainty and changes-of-mind are partially correlated, we used regression analysis to compare the time intervals over which the empirical and simulated CPP encoded these variables. The beta-coefficients plotted in Figure 4D show the effects of Certainty, changes-of-mind, and their interaction, on the real or simulated traces at each time-point. The empirical data showed a rising effect of Certainty on CPP amplitude starting ∼500ms before the initial response, which persisted until the confidence report. An effect of change-of-mind was also observed but it emerged much closer to the initial response and there was also a significant interaction between Certainty and change-of-mind that emerged prior to the initial response. Confirming the pattern shown by mean simulated DVs above, the Boundary-Single model was far better able to capture the time-course of the relationship between Certainty and amplitude than the Boundary-Distinct model. To give the Boundary-Distinct model a better chance of matching the pre-choice CPP effects, we ran additional versions of these models with the inclusion of a single drift rate variability parameter that was applied to both the pre- and post-choice drift rates. This allowed for the possibility that even a Distinct model could theoretically reproduce the CPP’s pre-choice scaling with confidence since trials with lower drift rates would be expected to result in less cumulative evidence at initial choice and final confidence report. However, the addition of drift rate variability resulted in numerically larger mean BIC for Boundary-Distinct (280 vs 279 without variability) and did not change the overall pattern of results, with Boundary-Single still providing a superior match to the CPP’s certainty effects and Boundary-Distinct still failing to capture the pre-choice amplitude effects (Extended Data Figure 4). The models were not readily distinguishable based on their ability to capture changes-of-mind, with Boundary-Distinct models again providing a better match to the pre-choice data but Boundary-Single models better capturing the post-choice data. However, the Boundary-Single model again substantially outperformed Boundary-Distinct in capturing the Certainty by change-of-mind interaction time-course.

As a final step we considered the possibility that a Single accumulation process could be implemented by two distinct signals, one that accumulates evidence for the initial choice and another that is initiated upon commitment and which tracks updates to the DV based on post-choice evidence (Boundary-Update model; see Methods and Extended Data Figure 5). This implementation provided a poor match to the CPP data, predicting larger pre- and post-choice amplitudes for trials with lower certainty ratings (the absolute DV in the Boundary-Update model will increase on average irrespective of whether the cumulative evidence favours the chosen or unchosen alternative), suggesting one continuous accumulation process explains the data better.

## Discussion

We adopted a neurally-informed modelling approach which enabled us to identify a set of computational mechanisms that could jointly account for choice behaviour, subsequent self-paced confidence reports, and the pre- and post-choice dynamics of a neural signature of evidence accumulation. The results indicate that rather than the accumulation process for confidence being subject to an internally set deadline on each trial, it is instead terminated once the cumulative evidence reaches a collapsing upper or lower confidence boundary. While the behavioural modelling unequivocally favoured a boundary-based stopping rule for the confidence reports, it was inconclusive on the question of whether evidence accumulation for confidence is simply a continuation of the process that informed the initial choice or whether it is the result of an independent accumulation process. Here, evaluating the degree to which the models could recapitulate neural data not included in the fitting process played a pivotal role in distinguishing between these alternative accounts, with the single process model markedly outperforming those invoking a distinct process in matching the CPP’s confidence-predictive dynamics before and after choices.

Theories of metacognition have long disagreed on the question of whether confidence is computed using the same computations as the choice, or whether a qualitatively different process of evidence accumulation informs confidence (4, 10). Here we examined this issue by pitting a model in which post-choice confidence reports arise from a continuation of the same evidence accumulation process that informed the initial choice (Boundary-Single) against models in which the accumulation process for confidence is driven by a separate process (Boundary-Distinct). Similar to the second-order model of Fleming & Daw (34), the Distinct models were blind to any variations in cumulative evidence at the time of the initial choice. This distinction between models could not be made in most previous studies since they have tended to assume fixed choice-boundaries, and therefore invariant cumulative evidence at the time of commitment to the initial choice (4, 6, 7, 34). Here, our model identified a substantial degree of boundary collapse for the initial choice process. As a result, later choices tended to be associated with less cumulative evidence and therefore lower accuracy, certainty, and confidence. While adding collapsing choice-boundaries did not produce a clear-cut winner between these models in the behavioural fits, it did have a significant impact when comparing the models based on their simulated evidence accumulation time-courses. Only the Boundary-Single model was able to reproduce the positive relationship between CPP amplitude and certainty that emerged several hundred milliseconds before the initial choice and remained until the final confidence report. In contrast, despite having the freedom to identify parameters that would allow for a distinct confidence-informing accumulation process to operate in parallel with the initial decision process, the Boundary-Distinct model nevertheless showed no pre-choice certainty effects and only small post-choice effects. Our modelling also considered the possibility of a post-choice accumulation process that is initiated at choice commitment, but which takes account of the evidence already accrued by adjusting its confidence boundaries (Boundary-Update). Here again, the model provided a poor match to the neural data, suggesting that not only is information shared between pre- and post-choice processes, but that it is encoded by a single continuous process rather than information transfer between two.

Our modelling results resolve an apparent divergence in previous empirical work regarding whether the computation of choice and confidence are serial or parallel processes. Much empirical work has shown how evidence presented after the choice causally affects subsequent confidence reports (16, 18, 35, 36) and yet here, as in several previous studies (12, 13, 37), neural correlates of confidence have been shown to emerge during decision-making. Although these observations are sometimes treated as opposing, they can all be accounted for within the context of the Boundary-Single model, in which a single evidence accumulation process informs the initial choice and continues past commitment in order to determine the final confidence judgment. The key reason why the Boundary-Single model is able to do so, is because it has collapsing choice boundaries such that already during the choice process some choices are made fast with strong evidence (and thus showing early traces of high confidence) whereas other choices are made slow with little evidence (and thus showing early traces of low confidence). Thus, a key insight from our modelling is that the neural processes governing metacognitive representations of choice confidence are intimately tied to the processes that produce the choices themselves. Our neurally-informed model furnishes new ways of disentangling choice and metacognitive processes by specifying pre- and post-choice boundaries, drift rate, and non-decision time parameters while also estimating the mapping of evidence to specific confidence ratings. In so doing, our model has the potential to offer a more detailed account of inter-individual differences in metacognitive performance including those associated with clinical brain disorders.

While our model points to a continuation of evidence accumulation after choice commitment, it is noteworthy that our participants nevertheless exhibited quite fast confidence-RTs on average. This is despite the fact that the physical evidence remained available on screen during the interval between the choice report and the confidence report deadline. For example, in the Accuracy Regime participants opted to report their confidence on average after 384ms, so over 2.5 seconds before the final deadline. Such short latencies for confidence RTs (compared to choice RTs which are often close to 1s on average) are not unique to our specific design, but are quite regularly observed in other studies as well (6, 7). The reasons for the limited duration of post-choice deliberation are unclear but one possibility is that participants may have implemented confidence-reporting boundary collapse rates that reflected the effort cost of sustaining attention over the relatively long trials durations implemented in this experiment (up to 3 seconds in the Speed Regime and up to 5.3 seconds in the Accuracy Regime).

There were however some features of the CPP that were not fully captured by our model, most notably its larger amplitude in the Speed Regime. This was an unexpected observation since previous studies that have applied speed-pressure to the *initial* choice have observed a lowering of pre-response CPP amplitudes (24, 29). One possible explanation is offered by the recent finding that asking participants to give post-decision confidence ratings resulted in slower and more accurate initial choices, along with a CPP that peaks earlier relative to the initial choice (11). This last observation suggests that post-choice confidence reporting may result in longer delays between initial choice commitment and response execution. Indeed, here too we observed that, on average, the CPP decayed back to baseline several hundred milliseconds prior to the final confidence report. It is possible that these delays are reduced and less variable in the Speed Regime resulting in a signal that is more closely time-locked to response execution and therefore larger in amplitude; correspondingly the Boundary-Single model identified a shorter post-choice non-decision time for non-change-of-mind trials in the Speed Regime. We did not fit variability parameters, to simplify the post-decision models, but future work could investigate whether the addition of non-decision time variability might enable the model to capture speed-pressure effects on the CPP.

Our behavioural and modelling results converge with those of a previous study using a similar task that separated speed-accuracy trade-offs (SAT) at the initial choice and confidence-report stages (6). We replicated their observation that increasing speed pressure for confidence reports impairs metacognitive sensitivity without decreasing mean confidence ratings, as well as decreasing choice RT. One striking difference is that Herregods et al. (6) observed significant confidence-SAT effects on the height of confidence-boundaries, whereas in the current work our confidence deadlines influenced the collapse rate of the confidence boundaries. This difference likely arises from different speed pressure manipulations; whereas we adjusted the response deadlines for the confidence reports, Herregods et al. (6) instead instructed participants to prioritise speed or accuracy, without specifying a new deadline. These two SAT manipulations have been shown to affect initial boundary height and collapse rates differentially (23) and the same may be true for confidence boundaries.

While we have described the Boundary-Single model in terms of a single accumulation process that continues seamlessly after initial choice commitment, this model variant did identify some notable differences in pre-versus post-choice parameter estimates. As mentioned, non-decision times were shorter for non-change-of-mind trials in the Speed Regime and the model also indicated that post-choice drift-rates were substantially larger than pre-choice drift-rates. Some previous studies that modelled pre- and post-choice behaviour have reported a lower post-decision drift-rates (38–40) which may arise from the fact that the physical evidence was extinguished once the choice was reported in those studies. Here, the same evidence that was presented before the choice remained on screen until confidence was reported and our modelling suggests that an increased weighting was applied to the post-choice evidence. However, Herregods et al. (6) reported higher post-choice drift rates despite also extinguishing the evidence after the initial choice, therefore more work is required to establish the origins of these effects, and a neurally informed approach will be useful in comparing mechanisms with similar behavioural predictions. Although not the focus of the current study, our model offers a useful framework for experimental manipulations probing post-choice changes to evidence sensitivity and phenomena such as confirmation bias (36, 41, 42).

## Methods

Prior to commencing any data analysis, the study design, hypotheses, and planned behavioural and neural analyses (with the exception of the modelling) were pre-registered on Open Science Framework (https://osf.io/adctm).

### Ethics

Trinity College Dublin ethics committee approved the study, and all procedures were performed in accordance with the Declaration of Helsinki and EU GDPR. Participants provided written consent before the start of the first session.

### Participants

In order to collect a sufficient number of change-of-mind trials per participant (which are usually around 10% of trials in similar tasks), we used a large-trial number, small-N design, collecting 2160 trials per person across three testing sessions. Participants were aged 18-40 without psychiatric or neurological illnesses. In our pre-registration we specified a sample-size of 12-15 participants based on previous studies using similar tasks (12, 43) but ultimately recruited 16 participants, as one participant had to be excluded prior to completing testing due to low accuracy during training, and another was excluded after indicating in their last testing session that they had misunderstood the instructions relating to the confidence ratings. The final dataset was therefore comprised of 14 participants (5 female & 9 male) with a mean age of 25 years (SD=4.3, range=20-36). They were paid €80-100 in total, including a performance-dependent bonus (see task details below).

### Design

We used a within-subject design to test the effect of varying the deadline (relative to the initial choice report) for making post-choice confidence reports. This was implemented with a blocked-design, alternating blocks of long and short deadlines, with the condition order counterbalanced across participants via pseudo-randomisation.

### Experimental Task

The task was a random-dot motion direction discrimination task designed in MATLAB (R2013b, www.mathworks.com) and Psychtoolbox-3 (Kleiner et al., 2007), and presented on a 1024 × 768 pixel monitor (100Hz refresh rate) approximately 60cm from the participant. Gaze and pupil diameter were recorded with an Eyelink 1000, following 9-point calibration at the start of each block, and drift-correction every 9 trials.

The task started with a central fixation cross and a white ring (10° diameter) on a grey background for 400ms, and then a cloud of 60 moving dots (0.15° diameter) appeared within an 8° diameter aperture. The dots initially moved with zero coherence for 800ms, to allow any visual-evoked potentials to reduce, before coherent motion to the left or right began (50% probability of each direction). After this lead-in period, coherent motion began with a proportion of dots being displaced relative to their position 3 frames earlier, to give a dot speed of 6° per second, with the remaining dots displaced randomly. To remove ambiguity about the onset of coherent motion, and reduce premature responses, a beep (100ms, 500Hz, Hanning-tapered at edges to prevent clicks) was presented at coherence onset. Participants responded with their left/right thumbs (on buttons ‘C’ and ‘N’) to indicate the perceived direction, with a deadline of 1500ms. After a valid response, the dots remained moving at the same coherence until participants reported their confidence in the perceived direction on a 6-point scale with their fingers (certain/probably/maybe left using keys 1, 2 and 3 respectively, or maybe/probably/certain right using keys 8, 9, and 0). Participants were instructed that they could change their minds between initial and confidence responses, for instance responding ‘left’ initially and then ‘maybe right’. We manipulated the deadline for this confidence-report, alternating blocks of trials with short (700ms, speed-pressure condition) or long (3000ms, accuracy-condition) deadlines while instructing the participants that the deadline for the initial choice always remained the same (1500ms). To continually remind participants which deadline applied on each trial, the fixation cross was coloured blue in long deadline blocks and yellow in short deadline blocks, and participants were also reminded of the change in deadline at the start of each block. The fixation cross changed colour at the end of a trial to indicate the accuracy of the initial response (green=correct, red=incorrect/early/late), and early or late initial-responses or late confidence-responses were given ‘TOO FAST’, ‘TOO LATE’ or ‘confidence TOO LATE’ feedback alongside a red fixation cross (feedback duration = 800ms). In each of 3 EEG sessions, participants completed 6 blocks of 120 trials, with a short break halfway through each block (2160 trials in total, 1080 per condition). Participants were reminded of the task instructions at the start of each session.

Prior to commencing the formal testing, a training session was conducted in which participants were first introduced to the different elements of the task and given time to practice and demonstrate to the researcher that they had understood the instructions. Training started out with 50% coherence until participants responded with >90% accuracy (initial-responses only, confidence-responses were not given here), and then coherence reduced in steps of 5% every 8 trials until it reached 10% coherence. Then the confidence-responses were introduced, and participants were given 30 trials to practice with the long (3000ms) deadline, and 30 trials with the short (700ms) deadline, using an easy coherence level based on their performance on the decreasing staircase. Following this, participants completed an additional block of 120 trials in which coherence was titrated using the QUEST procedure (44) to achieve 70% initial choice accuracy (participants were still asked to provide post-choice confidence ratings but these did not inform the titration procedure). A further 30 trials were run to verify that participants were performing within 65-75% accuracy. If performance was outside this range, then coherence was manually adjusted and further 30 trial blocks were run until accuracy was within the desired range. In addition to this initial staircase procedure, initial choice accuracy was monitored throughout the formal testing and the coherence level was manually adjusted if participants’ accuracy was not within 65-75%; this was only adjusted every second block, to ensure the coherence was the same for each pair of Speed and Accuracy blocks.

### EEG recording & pre-processing

EEG was recorded with a BioSemi ActiveTwo system (Biosemi, Netherlands) at 1024Hz with a 128-channel cap and two VEOG electrodes on the left eye. Data were pre-processed with ERPLab (45) and custom MATLAB scripts. Each block’s data were linearly detrended and low-pass filtered (40Hz FIR filter), and synchronised with the eye-tracking data with the EYE-EEG extension (46). EEG data were epoched from -1000:4000ms relative to coherence-motion onset, and baseline-corrected to the mean voltage in the 200ms before the coherence onset. Channels were visually inspected for extreme variance and up to 10% of channels interpolated.

Trials were examined for artefacts from the 50ms before the start of the baseline period until 250ms after the confidence-report on each trial. Trials were rejected if they contained scalp voltage ±100µV, bipolar VEOG voltage ±200µV, saccades over 2° or eye-position drifting >4° from the fixation-cross (as detected by Eyelink), or initial RT <100ms or no valid initial or confidence responses. The mean number of trials per participant without artefacts was 1807 (SD=230.5, range=1371-2063).

Voltages were transformed into Current Source Density (47) using the default parameters. This sharpens the spatial focus of components and reduces the overlap of frontal components that can obscure confidence-related effects from centro-parietal components (1).

Due to the auditory beep coinciding with coherent motion onset, there was a strong auditory evoked potential evidence in the EEG data, which we removed with the RIDE algorithm (48) similar to previous work (29). This algorithm separates out a stimulus-locked component from a response-locked component and a third component not locked to either stimulus or response times. These components are generated for each electrode from an entire session’s data and are fit over iterations to reduce the discrepancy with the data. Mean components are fit across each day’s session separately, to allow for slight variations in the auditory evoked potential on each day, but across the speed and accuracy conditions. This gave one stimulus-locked component per session that was the same for all trials and both conditions, and thus does not affect the differences in potentials between the two conditions. The stimulus-locked component was removed from all trials and electrodes, and the baseline correction re-applied.

The CPP was measured as the mean of 5 centro-parietal electrodes which showed a positivity in the grand-average topographies from -170:-50ms before the initial-response.

### Behavioural and Neural Analyses

Confidence was rated on a 6-point scale relative to the motion direction, which allowed for changes-of-mind between responses, and this was transformed to be relative to the initially-chosen direction, such that sticking with the same direction was rated as 4/5/6 for maybe/probably/certain non-change-of-mind, and change-of-mind responses were rated 3/2/1 for maybe/probably/certain change-of-mind. Thus, confidence measures participants’ subjective perception that their initial choice was correct. We additionally calculate ‘certainty’ in the final choice, using the maybe/probably/certain label selected (1/2/3), regardless of whether this was a change-of-mind or not. Initial-RT is measured in milliseconds from the coherent motion onset, and confidence-RT is measured in milliseconds from the time of the initial response (no minimum confidence-RT threshold is applied). RTs were log transformed for statistical analysis. We additionally calculate the area under the ROC2 curve (AUROC2) to measure metacognitive sensitivity (49); this measures how well subjective confidence-ratings track the objective accuracy of the initial choice.

To compare effects on behavioural variables, we used single-trial generalised linear mixed-effects regression for statistical analysis (using logistic regression for the binary variables of accuracy and change-of-mind). Predictor variables were z-scored to give standardised regression coefficients. The random-effects structures were picked using a forwards-selection method for each analysis separately (50), starting from a random intercept only and adding in terms in order of their BIC improvement until likelihood ratio tests became non-significant or the model failed to converge. Interaction-term random effects were only included if all the main-effects within them had already been added.

Additionally, we ran a window-free analysis to compare effects on the CPP across all time-points, by running the single-trial linear mixed-effects regressions at each time-point, as described above. The same random-effects structure was used for all time-points, which was the one chosen using the selection method described above for the mean-amplitude within the -150:-50ms time-window. To generate p-values, we ran 5000 permutations, permuting rows of the design matrix (i.e., the predictors) within-participants, and calculating the proportion of permutations that had a cluster mass as or more extreme than the true t-values (51, 52), to give a two-tailed p-value with family-wise error rate controlled at .05. These permutation-tests are time- and computationally-intensive across many time-points and trials due to mixed-effects regressions being much slower than fixed-effects methods, so we used the lmeEEG toolbox (53) to speed this up. This method first regresses out the random-effects at every time-point, by fitting the mixed-effects model, removing the random-effects, leaving behind the fixed-effects and residuals. This ‘marginal EEG’ data is then analysed with fixed-effects regressions using OLS (for both true-regressions and permutation-regressions), which gives almost identical results to running the full mixed-effects permutation testing, but much quicker (53).

We set a minimum trial-number threshold for inclusion in the figures/analyses of 10 trials per condition (or combination of conditions as in the example above). Participants who had fewer than 10 trials per condition/combination were excluded from that particular condition, but included in all other conditions and analyses. Please note, the mixed-effects analyses used allow missing data for particular conditions.

When plotting CPP traces, the data were smoothed with a 100ms window moving-average.

### Model Specification

We modelled the initial decision using a drift-diffusion process with linearly collapsing boundaries. Here, noisy evidence accumulation is approximated as a discrete-time random walk:

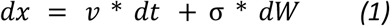

where *dx* represents the change in evidence over the time step *dt* (set to 0.001s), *v* is the drift rate representing the average rate of evidence accumulation, and σ represents the within-trial diffusion noise, which was fixed to 1. *dW* represents an increment of a Wiener process, a random variable drawn from a normal distribution with a mean of 0 and a variance of dt. The starting point of the accumulation process was set to *z* * *a*, where *z* was fixed to an unbiased starting point of 0.5, and *a* represents the upper decision boundary (with 0 representing the lower boundary). To account for the observed RT distributions, which showed a low degree of skew and a strongly decreasing conditional accuracy function, likely induced by the sharp response deadlines, we incorporated a linear boundary collapse mechanism, controlled by an urgency parameter, *u* (model comparisons of initial-decision fits confirmed that adding *u*, and fixing *z*, improved the model fits via lower BIC). The upper boundary at time *t* was defined as *a(t) = a - u*t*, and the lower boundary was defined as *a(t) = 0 + u*t*. A choice and corresponding RT were registered when the accumulated evidence crossed either the upper or lower boundary. We used an accuracy-coding scheme, such that crossing the upper boundary corresponded to a correct response and crossing the lower boundary corresponded to an incorrect response. Finally, a non-decision time parameter, *ter*, was added to the predicted RT to account for processes unrelated to the decision itself, such as stimulus encoding and motor execution.

To inform the calculation of subsequent confidence judgments, an evidence accumulation process for confidence was instantiated. The accumulation process for confidence was modelled with a drift rate (*v2*) that was allowed to differ from the drift rate for choices (*v*) (39). The accumulation for confidence continued until a specific criterion was met, with the nature of this criterion differing across the different classes of models we investigated. We evaluated four classes of models, distinguished along two key dimensions: (1) Time-based and Boundary-based models were compared to identify the stopping rule governing the termination of accumulation for confidence and (2) Single vs. Distinct models were compared to determine whether the accumulation process for confidence operates within the same reference frame as the initial choice (i.e. evidence for left versus right motion) or restarts in a new reference frame (e.g. accumulating evidence for correct vs error). Note that within the single process models the accumulation for confidence can be thought of as post-decision evidence accumulation (4) whereas for the distinct models given that the accumulation process for confidence is distinct from that for choices these models are agnostic with respect to the question whether the accumulation process is post-decisional or whether they run in parallel with the accumulation for choices.

Regarding the stopping rule, in time-based models, accumulation for confidence terminates after a certain duration. This duration can be fixed, such as during the non-decision time (e.g., “pipeline evidence”, (3)), set equal to the average observed time of confidence responses, or simply estimated freely (2DSD; (4)). In Boundary-based models, accumulation for confidence terminates when the accumulated evidence reaches a collapsing confidence boundary (6, 7). Regarding the reference frame, Single-process models assume that evidence accumulation for confidence continues within the same reference frame as the initial decision (12). Consequently, the starting point for the confidence judgment is determined by the end point of the initial decision process. Distinct-process models, however, posit that evidence accumulation for confidence is initiated with a new reference frame independent of the initial decision’s starting and end points (18). This allows for the possibility that confidence formation could, in principle, run partly in parallel with the initial decision. Another crucial difference with respect to the different reference frames is how differential evidence is translated into absolute evidence, which is further discussed below, in the section on simulating the decision variable.

All models included the following parameters: a drift rate for confidence (*v2*), two non-decision time components (*ter2*_*no-CoM*_, *ter2*_*CoM*_) to account for differences between change-of-mind and non-change-of-mind trials (as changes of mind required a hand switch), and five confidence criteria (*c1* to *c5*). For convenience, during fitting on incorrect trials the post-decision drift rate (*v2*) was multiplied by -1, reversing the direction of accumulation, to reflect the fact that evidence on error trials predominantly supports the incorrect response, which supports metacognitive sensitivity. Note that this is equivalent to keeping the drift rate the same, but flipping the confidence mapping on those trials (i.e., computing confidence conditional on the response).

In the Time-based models, evidence accumulated until a deadline (*τ*) was reached. Normally distributed noise (*σ*_*τ*_) around this deadline was included to account for variability in the termination time. In the Boundary-based models, evidence accumulated until it reached one of two collapsing confidence boundaries. For Distinct process models, evidence accumulation started at a new, freely estimated starting point (*z2*) and continued until the timer finished (Time-Distinct model) or it reached either the upper or lower collapsing confidence boundary (Boundary-Distinct model; defined by boundary separation parameter *a2*, and two separate collapse rate parameters, *u2*_*up*_, *u2*_*down*_). We allowed for negative non-decision times in the Distinct models, consistent with the idea that the initial decision and confidence formation processes can overlap in time. For Single process models, evidence accumulation started at the point where the initial decision boundary was crossed and continued until the timer finished (Time-Single model) or a new set of collapsing confidence boundaries were reached (Boundary-Single model). These confidence boundaries were defined by parameters *a2*_*up*_ and *a2*_*down*_ at time 0, and their collapse was controlled by separate urgency parameters (*u2*_*up*_, *u2*_*down*_).

Finally, the accumulated evidence at the termination of the post-decisional process was mapped onto a discrete six-point confidence rating. This mapping involved adding metacognitive noise (with standard deviation *σ*_*meta*_) to the final evidence value, representing noise in the readout process (54, 55). The resulting value was then categorized into one of six confidence levels using five estimated confidence criteria (*c1* to *c5*). To enforce monotonicity, we estimated the first criterion (c1) and the non-negative differences between successive criteria, rather than fitting all five independently. The six confidence levels correspond to “certain”, “probably”, “maybe” responses for change-of-mind (ratings 1, 2, and 3) and “maybe”, “probably”, “certain” non-change-of-mind trials (ratings 4, 5, and 6).

### Model Fitting Procedure

We employed a two-stage fitting procedure to estimate the parameters of the four confidence models. This approach ensured that all models were based on the same pre-decisional mechanisms, allowing us to isolate the effects of different processes on confidence formation (simulations confirmed the two-stage fitting could accurately recover models and parameters, see details below and Extended Data Figure 6). In the first stage, we fit the initial decision data using a drift-diffusion model (DDM) with linearly collapsing boundaries. To find the best-fitting parameters for this initial decision model, we minimized a goodness-of-fit metric (*G*2) that compared the observed and model-predicted RT distributions. Specifically, we calculated the *G2* likelihood-ratio statistic as follows:

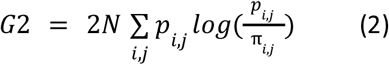

where *N* is the total number of observed trials, *p*_*i,j*_ and *π*_*i,j*_ are the observed and simulated proportions of RTs, respectively, that fall within a given quantile bin *i* for each response *j* (correct or incorrect). We used the following quantiles: 0.1, 0.3, 0.5, 0.7, and 0.9, creating 5 bins for correct responses and 5 bins for incorrect responses. The simulated proportions were obtained by simulating the DDM 5000 times for each iteration of the optimization routine. We used Differential Evolution (DEoptim package in R, (56)) to perform the parameter optimization, running the algorithm for 500 iterations. This procedure was performed separately for each participant and speed condition. The best-fitting parameters (*v, a, ter, u*) from this initial decision model were then fixed in the second stage when fitting the confidence models.

In the second stage, we fit the four classes of confidence models to the confidence ratings and confidence-RTs. The parameters of the models (*v2, ter2*_*no-CoM*_, *ter2*_*CoM*_, *τ, σ*_*τ*_, *a2, z2, u2*_*up*_, *u2*_*down*_, *a2*_*up*_, *a2*_*lower*_, *c1* to *c5, σ*_*meta*_) were estimated by minimizing a composite objective function. This combined three equally weighted goodness-of-fit metrics, each designed to capture a distinct aspect of the behavioural data. Each of the metrics was in the form of a likelihood-ratio statistic, G2, and calculated as specified in formula 2. However, the specific meaning of the indices *i* and *j* varied depending on the metric. A first metric targeted the distribution of confidence-RTs conditional on decision accuracy. Here, *i* indexed the confidence-RT bins (created by the same quantiles as specified above) while *j* indexed the accuracy of the decision. Second, we used the overall proportion of responses for each of the confidence ratings conditional on decision accuracy. Here, *i* indexed the confidence levels (1-6) while *j* indexed the accuracy of the decision. Finally, we calculated the proportion of responses for each confidence rating conditional on five equally-spaced confidence-RT bins to capture the relationship between confidence and confidence-RT. Here, *i* indexed the confidence levels (1-6), while *j* indexed the confidence-RT bin (1-5). The three *G*2 values were summed to create a composite goodness-of-fit metric that was minimized. We opted for equal weighting of the metrics as we had no a priori reason to prioritize one aspect of the data over another. This approach, using multiple complementary goodness-of-fit metrics, allowed us to constrain the models with different aspects of the behavioural data, ensuring a robust estimation of the post-decision processes.

To compare the fit of the four model classes, we computed the Bayesian Information Criterion (BIC) for each model and each participant, according to the following formula:

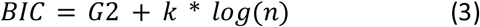

where k is the number of free parameters, and n is the number of trials (57). Lower BIC values indicate a better fit while taking into account model complexity. In addition to these quantitative metrics, we visually inspected the model fits to assess how well each model captured the key patterns in the behavioural data, including the shapes of the confidence-RT distributions, the relationship between confidence and confidence-RT, and the effects of speed pressure on choice and confidence (see Results and Figure 3).

We validated our model and parameter recovery procedures by simulating data for each set of fitted parameters (1000 trials simulated per 14 participants and 2 conditions) for each of the four models (giving 112 simulations in total), and then fitting each of the four models to these simulated data. The ground-truth model was correctly identified via lowest BIC in 97/112 of these simulations, including 25/28 for the Boundary-Distinct and 28/28 for Boundary-Single. We then took the parameters originally fit to the true data, and correlated them with the parameters recovered from fitting the ground-truth model to simulated data. In general, for each model the recovered parameters highly correlated with the original parameters (r > .9), with only lower recovery for some for non-central parameters (e.g. 1st confidence criteria and urgency/collapse-rate for lower confidence boundary in some models). The full model and parameter recovery results can be found in Extended Data Figure 6.

### Simulated Decision Variable

To compare the latent decision variable from our models with the neural data (specifically the CPP), we simulated the decision variable’s trajectory over time, incorporating additional assumptions and noise sources to better reflect the variability observed in noisy neural scalp recordings and the characteristics of the CPP (Extended Data Figure 2). The decision variable was simulated in epochs ranging from -1s to +3s relative to stimulus onset, with a time step (*dt*) of 0.01s, equivalent to a 100 Hz sampling rate. We simulated 1000 trials for each subject and speed pressure condition. For each trial, we first simulated a pre-stimulus period and an initial non-decision period, assumed to reflect stimulus encoding. During this period, the decision variable was modelled as a noisy mean-reverting Ornstein-Uhlenbeck process (to prevent the DV from hitting the decision boundaries before the evidence began):

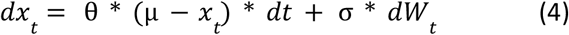

where *x*_t_ represents the value of the decision variable at time *t*, θ is the mean reversion rate (set to 10), *μ* is the long-term mean to which the process reverts (set to the starting point, 0), *σ* scales the noise level (set to 0.1), and d*W*_t_ is an increment of a Wiener process, drawn from a normal distribution with a mean of 0 and a variance of *dt*. The duration of the stimulus encoding stage was determined by subtracting a simulated motor execution time from the total non-decision time (*Ter*) estimated in the initial decision model. The motor execution time was assumed to be 0.1s on average, based on previous work (Kelly et al., 2021), and was drawn for each trial from a uniform distribution ranging +-0.15 s around the expected motor execution time.

Following the encoding stage, the initial decision stage started. Here, differential evidence accumulated according to a standard drift-diffusion process, with drift rate *v* (formula 1). The decision boundaries were transformed so that the starting point of accumulation was 0, rather than *a*\*z. The boundaries were linearly collapsing, as determined by the urgency parameter *u* from the initial decision model. To approximate the CPP, we computed the absolute value of accumulated differential evidence, i.e., reflecting evidence around zero. This approach is motivated by empirical findings showing that the CPP displays a positive-going increase regardless of the ultimate decision, and is hypothesized to represent the summed activity of two distinct neural populations. Each of these populations encodes evidence for a specific choice option, both with positive polarity, unlike the differential evidence representation typically used in the DDM (24).

The nature of the post-decisional stage depended on the specific process type (Distinct or Single). In Distinct models, a separate evidence accumulation process was initiated with its own starting point. The accumulated evidence in this stage was reflected around this new starting point (*z2*) to represent the absolute value in the reference frame of confidence formation. The simulated trajectories of the initial and post-decisional processes were then summed to generate the final decision variable trajectory. Conversely, Single-process models assumed a continuous accumulation of evidence from the initial decision stage, without a shift in the reflection point. The absolute evidence in these models was calculated throughout the trial by reflecting the accumulated evidence around the original starting point (*z*). Thus, the critical distinction between the Distinct and Single-processes lies in whether the reference frame for evidence accumulation remains fixed throughout the trial and is specified in terms of the original decision (Single) or undergoes a change to form a new, independent process for post-decisional evaluation (Distinct).

The Boundary-Update model used the same fitted parameters as the Boundary-Single model and differed only in terms of how the post-choice accumulation process was simulated. Instead of being a continuation of the accumulation process that determined the initial choice, in Boundary-Update the pre- and post-choice stages of accumulation were simulated with separate signals, with the post-choice signal onsetting at choice-boundary-crossing and the pre-choice process decaying to zero. The post-choice process accumulated post-decision evidence until it crossed a confidence-boundary. This allowed initial-boundary crossing values to affect how close to the confidence boundaries the post-decision process began Following the post-decisional stage, the level of confidence was determined for all models using the mechanisms described previously. Subsequently, a second non-decision time stage was initiated, representing the time required to execute the confidence response. The duration of this stage was sampled from a uniform distribution centred on the model’s fitted mean confidence non-decision time, with a range of ±0.15 s. Separate means were used for change-of-mind trials (*ter2*_*CoM*_) and non-change-of-mind trials (*ter2*_*no-CoM*_), to account for the additional motor time required due to changing the response hand. On change-of-mind trials, evidence accumulation was allowed to continue during this confidence non-decision time stage, consistent with the hypothesis that pipeline evidence (i.e., evidence received during the non-decision processes) is continuously integrated even after the initial decision (3). On non-change-of-mind trials, or upon completion of the confidence non-decision time stage on change-of-mind trials, the accumulated evidence was allowed to decay back towards the starting point. This decay was modelled as a noisy linear decay process:

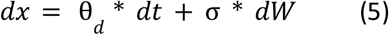

where *θ*_*d*_ represents the linear decay factor (set to -5; the results were similar for other decay rates). The noise scaling parameter, σ, was the same as that used in the pre-stimulus stage. This decay process ensured that the simulated decision variable gradually returned after the initial response (for very short post-decisional accumulation times) or confidence response had been executed. When plotting the simulated decision variable, the data were smoothed with a 100ms window moving-average (cf. the CPP).

## Data Availability

Anonymous data are publicly available on OSF (https://osf.io/4bp96/).

## Code Availability

MATLAB code for all analyses is available on GitHub (https://doi.org/10.5281/zenodo.15586067) and the R code for modelling is available on OSF (https://osf.io/f6xyw/).

## Acknowledgements

This study was supported by a grant from Horizon 2020 European Research Council Consolidator Grant IndDecision 865474, and grants from the Research Foundation Flanders (1242924N) awarded to L.V., and (G0B0521N) awarded to K.D.

## Author contributions

JPG: Conceptualisation, Methodology, Software, Validation, Formal analysis, Investigation, Data curation, Writing-Original Draft, Writing-Review & Editing, Visualisation, Project administration.

LV: Conceptualisation, Methodology, Software, Validation, Formal analysis, Investigation, Writing-Original Draft, Writing-Review & Editing, Visualisation.

SLM: Investigation, Writing-Review & Editing.

CMC: Investigation, Writing-Review & Editing.

DM: Investigation, Data curation, Writing-Review & Editing.

KD: Conceptualisation, Methodology, Resources, Writing-review & editing, Supervision, Funding Acquisition.

ROG: Conceptualisation, Methodology, Resources, Writing-review & editing, Supervision, Funding Acquisition.

