## Extended Data for "A single evidence accumulation process informs perceptual choices and subsequent confidence reports"

### Supplementary Information

#### Extended Data Tables

***Extended Data Table 1. Pre-decisional parameters for the initial-choice model (all four post-decision models used the same initial-choice parameters), in speed and accuracy conditions.*** *This model gave better fits than a model without urgency (u; ΔBIC = 75).*

| Model | Parameter | Speed | Accuracy | *t* | *p* | Sig. |
|---|---|---|---|---|---|---|
| Pre-decision | *a* | 1.9915 | 2.1655 | 2.4849 | 0.0274 | * |
| | *ter* | 0.3774 | 0.3861 | 0.8631 | 0.4038 | n.s. |
| | *u* | 0.8897 | 0.9619 | 1.726 | 0.108 | n.s. |
| | *v* | 0.7513 | 0.7505 | -0.035 | 0.9726 | n.s. |

*Note. v = drift rate, a = decision boundary, ter = non-decision time, u = urgency rate. Degrees of freedom for the t-test is 13.*
**=p<.05, **=p<.01, ***=p<.001, ****=p<.0001.*

*Extended Data Table 2. Post-decisional parameters for the four models, in speed and accuracy conditions.*

| Model | Parameter | Speed | Accuracy | *t* | *p* | Sig. |
|---|---|---|---|---|---|---|
| Time-Distinct | $\sigma_{meta}$ | 2.5126 | 2.9097 | 3.7856 | 0.0023 | ** |
| | $\sigma_{\tau}$ | 0.0781 | 0.1544 | 3.2583 | 0.0062 | ** |
| | $\tau$ | 0.4114 | 0.4932 | 1.8468 | 0.0877 | n.s. |
| | *ter2A* | -0.2082 | -0.1548 | 0.9132 | 0.3777 | n.s. |
| | *ter2B* | 0.1011 | 0.2497 | 1.5785 | 0.1385 | n.s. |
| | *v2* | 0.461 | 0.7854 | 2.4647 | 0.0284 | * |
| Time-Single | $\sigma_{meta}$ | 0.5033 | 0.5467 | 0.666 | 0.517 | n.s. |
| | $\sigma_{\tau}$ | 0.078 | 0.1494 | 3.3215 | 0.0055 | ** |
| | $\tau$ | 0.1687 | 0.2815 | 2.9654 | 0.0109 | * |
| | *ter2A* | 0.0353 | 0.0616 | 2.2316 | 0.0439 | * |
| | *ter2B* | 0.3539 | 0.3679 | 0.1727 | 0.8656 | n.s. |
| | *v2* | 0.8188 | 0.9306 | 1.3047 | 0.2146 | n.s. |
| Boundary-Distinct | *a2* | 4.2447 | 4.2029 | -0.2222 | 0.8276 | n.s. |
| | $\sigma_{meta}$ | 1.2046 | 1.2898 | 0.5515 | 0.5906 | n.s. |
| | $ter2_{no\text{-}CoM}$ | 0.0027 | -0.0121 | -0.2896 | 0.7767 | n.s. |
| | $ter2_{CoM}$ | 0.2414 | 0.2134 | -0.2499 | 0.8065 | n.s. |
| | $u2_{down}$ | 2.6197 | 2.5474 | -0.1431 | 0.8884 | n.s. |
| | $u2_{up}$ | 4.1849 | 2.6221 | -4.2081 | 0.001 | ** |
| | *v2* | 0.5332 | 0.715 | 1.8077 | 0.0938 | n.s. |
| | *z2* | 0.7789 | 0.7425 | -0.7087 | 0.491 | n.s. |
| Boundary-Single | $a2_{down}$ | 0.4383 | 0.3208 | -1.3315 | 0.2059 | n.s. |
| | $a2_{up}$ | 0.5251 | 0.4144 | -1.5322 | 0.1494 | n.s. |
| | $\sigma_{meta}$ | 0.908 | 0.9341 | 0.579 | 0.5725 | n.s. |
| | $ter2_{no\text{-}CoM}$ | 0.0357 | 0.0791 | 2.3082 | 0.0381 | * |
| | $ter2_{CoM}$ | 0.2546 | 0.2973 | 0.6833 | 0.5064 | n.s. |

| | | | | | |
|---|---|---|---|---|---|
| $u2_{down}$ | 5.9565 | 4.3347 | -2.1238 | 0.0535 | n.s. |
| $u2_{up}$ | 3.4699 | 1.4761 | -5.1554 | 0.0002 | *** |
| *v2* | 1.687 | 1.6808 | -0.0217 | 0.983 | n.s. |

*Note. v2 = confidence drift rate, z2 = confidence starting point, a2 = confidence boundary, ter2 = confidence non-decision time, u2 = confidence urgency rate, $\boldsymbol{\sigma}_{meta}$ = metacognitive noise, τ = average deadline, $\boldsymbol{\sigma}_{\tau}$ = variability in average deadline. If applicable, up/down refers to upper or lower boundary, Com/no-CoM refers to Change of Mind and no-Change of Mind. Degrees of freedom for the t-test is 13. *=p<.05, **=p<.01, ***=p<.001, ****=p<.0001.*

*Extended Data Table 3. Confidence criteria parameters for the four models, in speed and accuracy conditions.*

| Model | Parameter | Speed | Accuracy | *t* | *p* | Sig. |
|---|---|---|---|---|---|---|
| Time-Distinct | *c1* | -4.4907 | -4.5541 | -0.601 | 0.5582 | n.s. |
| | *c2* | 1.1086 | 1.1102 | 0.0171 | 0.9866 | n.s. |
| | *c3* | 0.6474 | 0.6237 | -0.3117 | 0.7602 | n.s. |
| | *c4* | 2.1571 | 2.1094 | -0.2201 | 0.8292 | n.s. |
| | *c5* | 2.4064 | 2.3914 | -0.1171 | 0.9086 | n.s. |
| Time-Single | *c1* | 0.1979 | 0.1874 | -0.1259 | 0.9018 | n.s. |
| | *c2* | 0.2851 | 0.3434 | 1.8105 | 0.0934 | n.s. |
| | *c3* | 0.1619 | 0.1776 | 0.7098 | 0.4904 | n.s. |
| | *c4* | 0.5432 | 0.5572 | 0.4017 | 0.6945 | n.s. |
| | *c5* | 0.571 | 0.6202 | 1.0357 | 0.3192 | n.s. |
| Boundary-Distinct | *c1* | 0.3777 | 0.3045 | -0.9392 | 0.3647 | n.s. |
| | *c2* | 0.5567 | 0.5578 | 0.0167 | 0.9869 | n.s. |
| | *c3* | 0.3246 | 0.303 | -0.3328 | 0.7446 | n.s. |
| | *c4* | 1.215 | 1.0086 | -1.3531 | 0.1991 | n.s. |
| | *c5* | 1.2327 | 1.266 | 0.296 | 0.7719 | n.s. |
| Boundary-Single | *c1* | 0.1339 | 0.0891 | -0.6077 | 0.5539 | n.s. |
| | *c2* | 0.2971 | 0.3703 | 1.7623 | 0.1015 | n.s. |
| | *c3* | 0.2205 | 0.2396 | 0.5071 | 0.6206 | n.s. |
| | *c4* | 0.8928 | 0.8317 | -0.7329 | 0.4766 | n.s. |
| | *c5* | 0.8914 | 0.879 | -0.2049 | 0.8408 | n.s. |

*Note. Degrees of freedom for the t-test is 13. *=p<.05, **=p<.01, ***=p<.001, ****=p<.0001.*

### Extended Data Figures

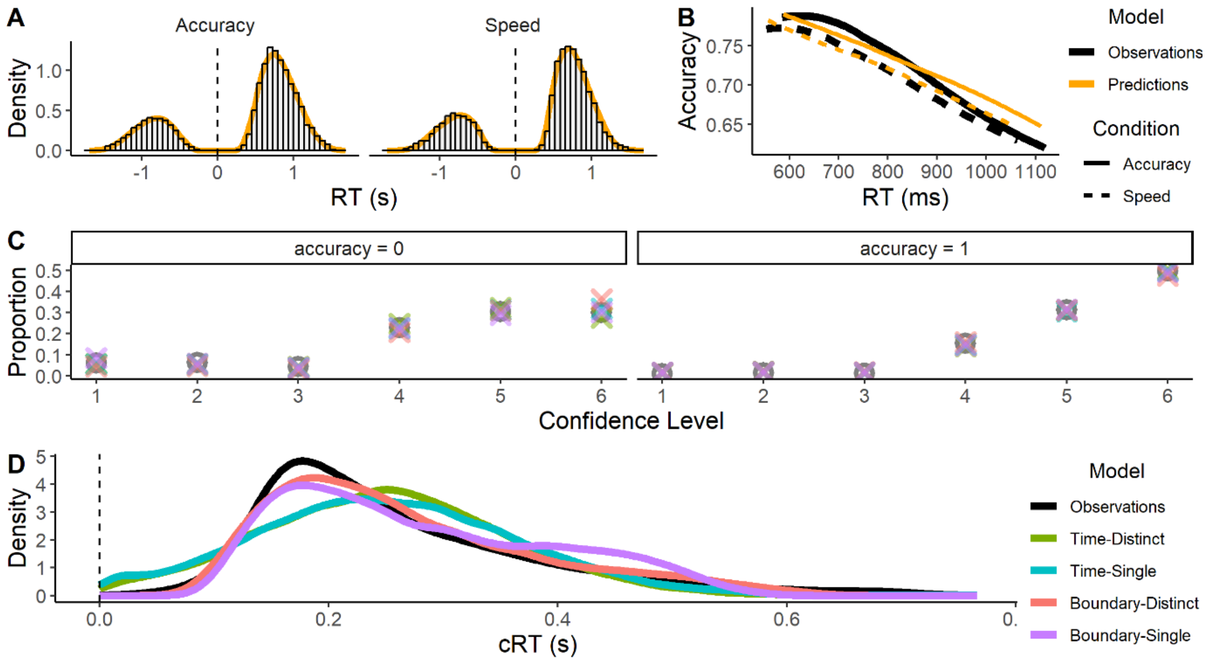


***Extended Data Figure 1. Additional model fitting data.*** *(A) Distributions of initial RTs for correct (positive) and incorrect (negative) choices in the Accuracy (left) and Speed (right) conditions. Histograms show observed data, while orange lines show model predictions. (B) Conditional accuracy functions showing the proportion of correct responses as a function of initial RT in the Accuracy (solid lines) and Speed (dashed lines) conditions. Black lines represent observed data, and orange lines represent model predictions. (C) Observed and fitted Confidence Frequencies conditional on decision accuracy. The Boundary-Distinct model (orange) was less precise at capturing these frequencies conditional on decision accuracy. (D) Confidence-RT (cRT) Distributions for a single example subject in the speed condition, showing that time-based models cannot capture the confidence-RT distributions when they are sufficiently skewed, leading to a worse overall fit in terms of confidence-RT.*

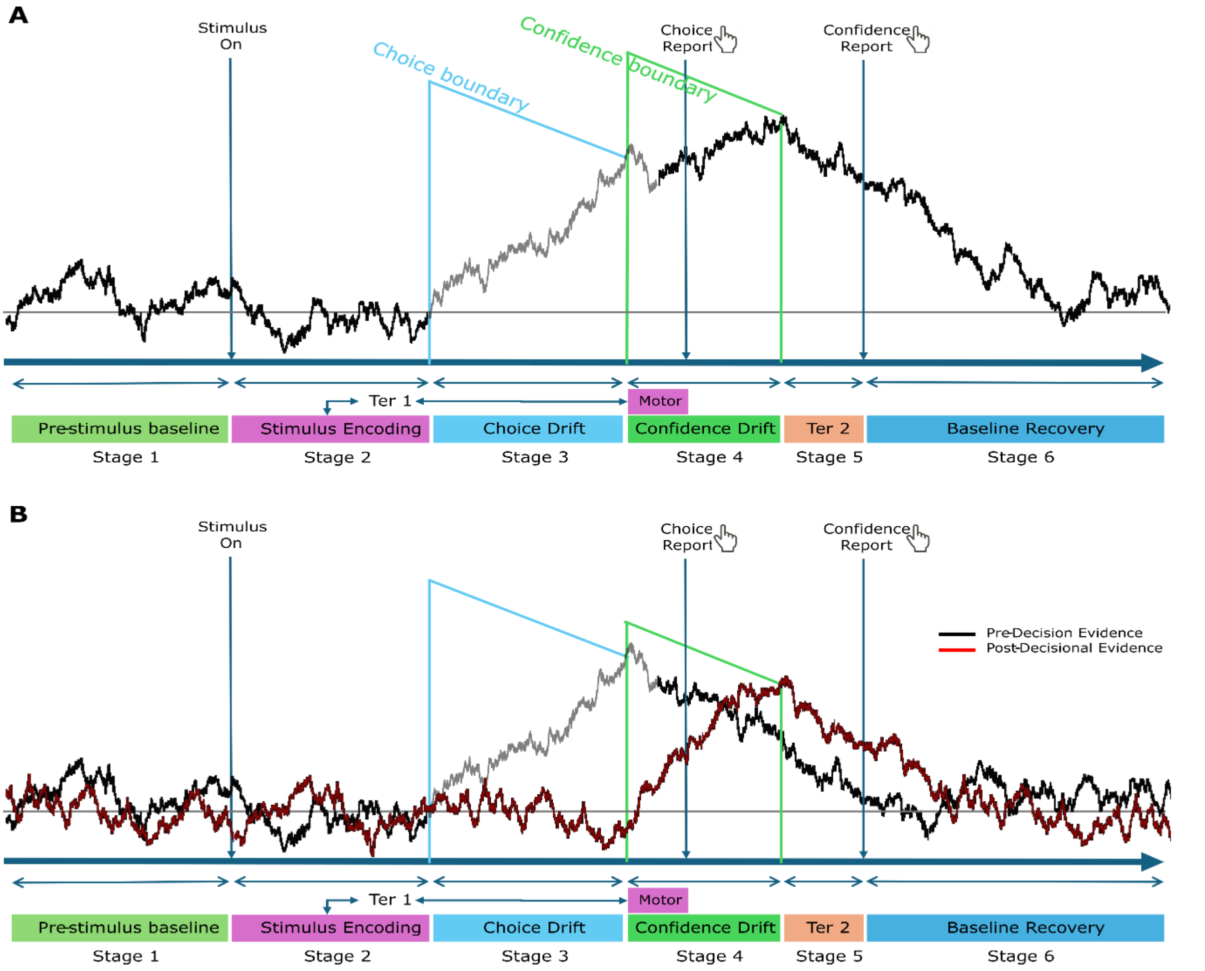


***Extended Data Figure 2. Stages of the simulated decision variable for boundary-based models.*** *(A) Single-Process Model. (B) Distinct-Process Model. Single-process models have one accumulator that continues from initial-decision to confidence-accumulation, whereas the Distinct-process models have two accumulators, with the evidence switching from one to the other at the initial decision time. The sum of absolute decision variables was taken as the CPP proxy. When no evidence is given (before stimulus onset or after initial/confidence decision), the accumulators receive no evidence, solely accumulating noise, with a mean-reverting decay. Non-decision time is composed of sensory-encoding delay and motor execution delay, the latter was set to be 100ms with uniform variability of ±150ms, which was subtracted from the fitted non-decision time to calculate sensory encoding delay (Stage 2). After stimulus onset, noisy accumulation of evidence began (Stage 3) until it reached a linearly collapsing boundary (blue line; the lower-boundary corresponding to an error response is not shown), with motor-execution delay added to this time to give initial RT. The confidence-accumulation (Stage 4) starts upon initial-decision commitment; in the Single-process models this is the same accumulator as the initial one, whereas in the Distinct-process models, it is a separate process that only receives evidence at this point, having been accumulating noise with a decay until now. Confidence-accumulation continued until it reached a linearly collapsing confidence boundary (green line), with confidence non-decision time (Ter2) added to generate confidence-RT. After an accumulation process ends, the signal has a noisy linear decay.*

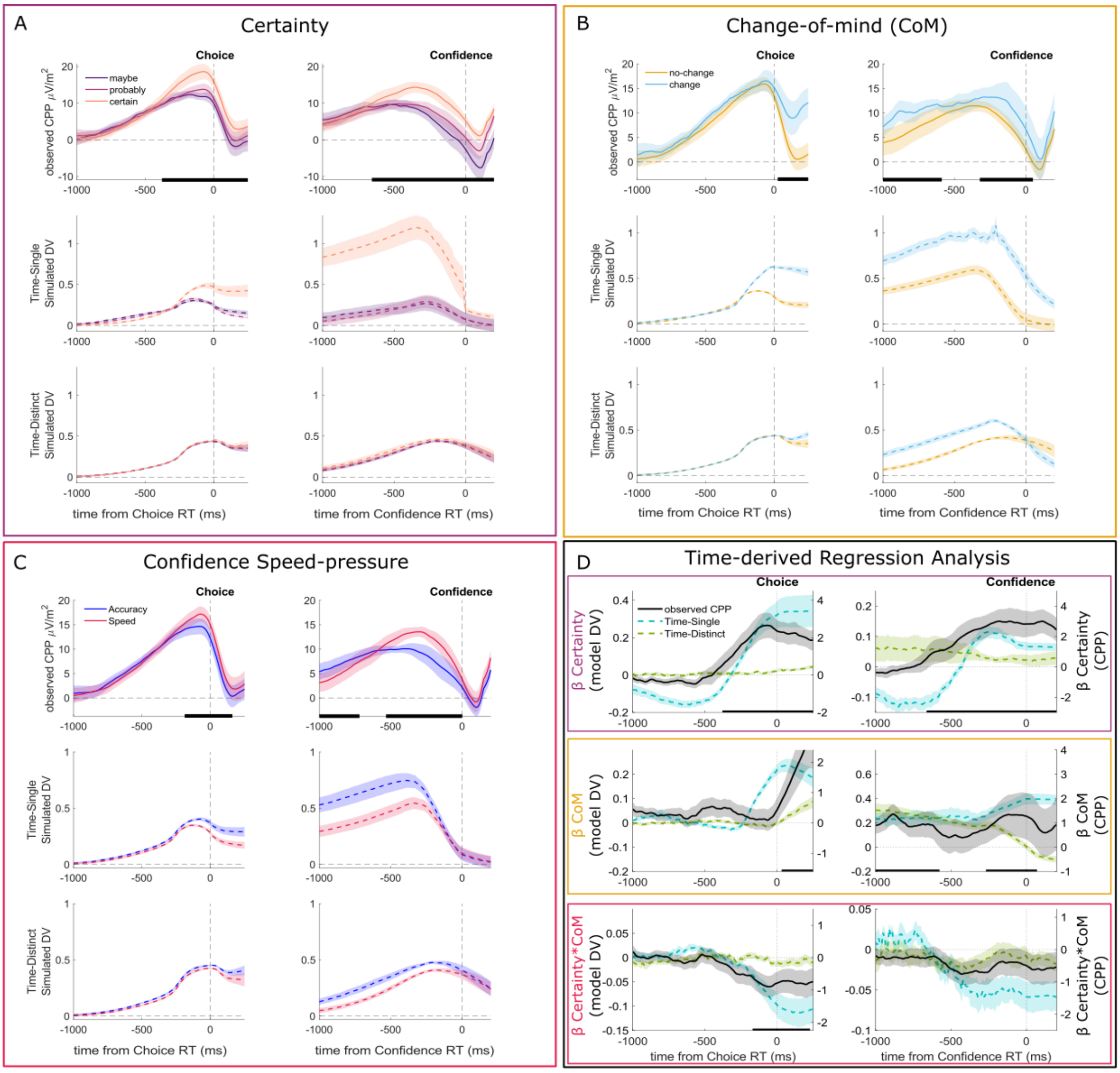


***Extended Data Figure 3. Simulated Decision Variable for Time-based models.*** *Observed CPP (solid lines) are the same data shown in main Figure 4. A) Certainty effects, B) Change-of-mind (CoM) effects, C) Speed-pressure effects. D) Time-based models regression analyses, running the same regression analyses as Figure 4D in the Time-Based models' absolute Decision Variables. Neither model can capture the qualitative effects fully; the Certainty effect is diminished for Time-Single and non-existent for Time-Distinct, while the CoM effect is positive before the initial choice for Time-Single and much too small in Time-Distinct. Additionally, both models predict positive effects of Speed-pressure, like the Boundary models.*

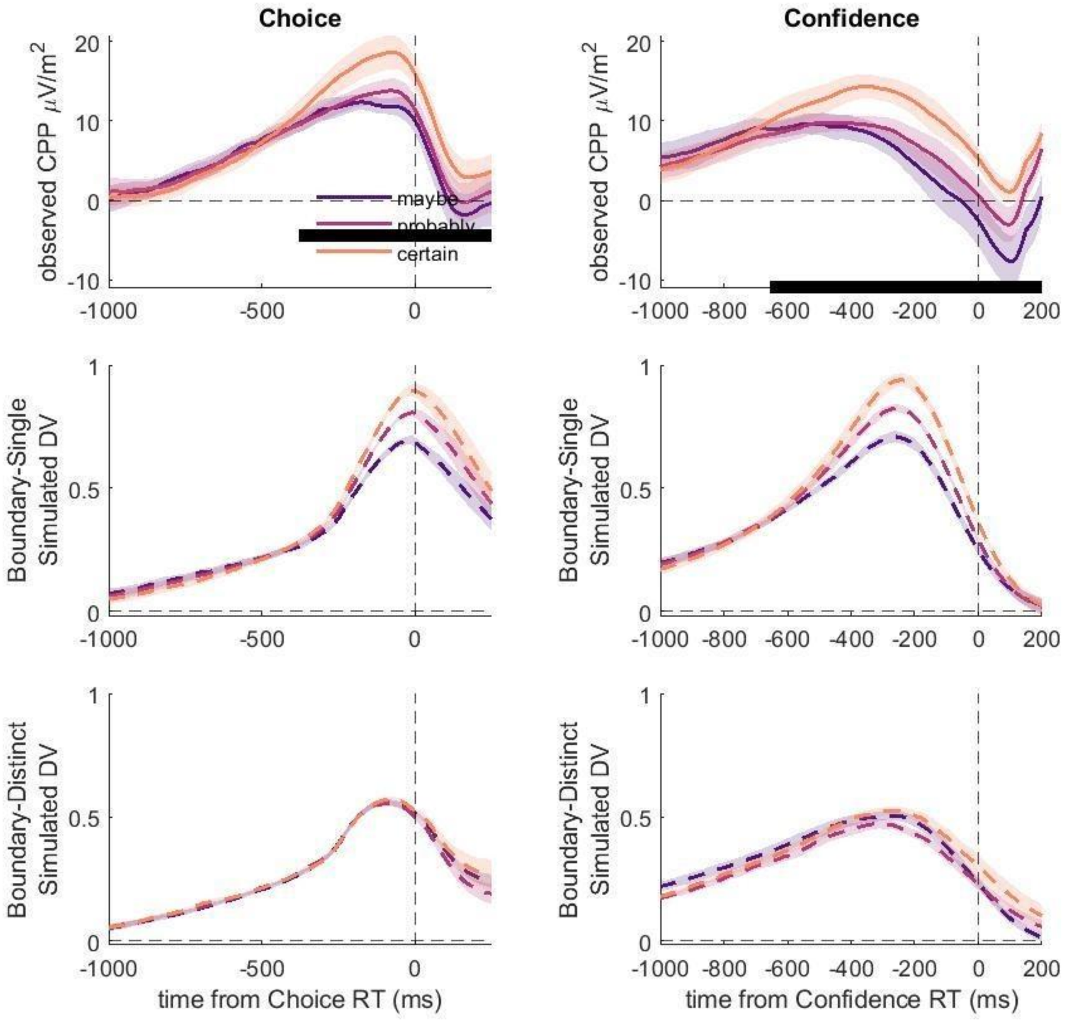


*Supplementary Figure showing simulated DVs for Boundary-Distinct and Boundary-Single for models with drift-rate variability. They are almost identical to the models without drift-rate variability.*

***Extended Data Figure 4. Simulated Decision Variable for Boundary-Distinct and Boundary-Single models with drift rate variability included.*** *The Boundary-Distinct model still failed to capture the observed certainty effects on the CPP even with the addition of drift rate variability that was shared across the pre- and post-choice accumulation phases.*

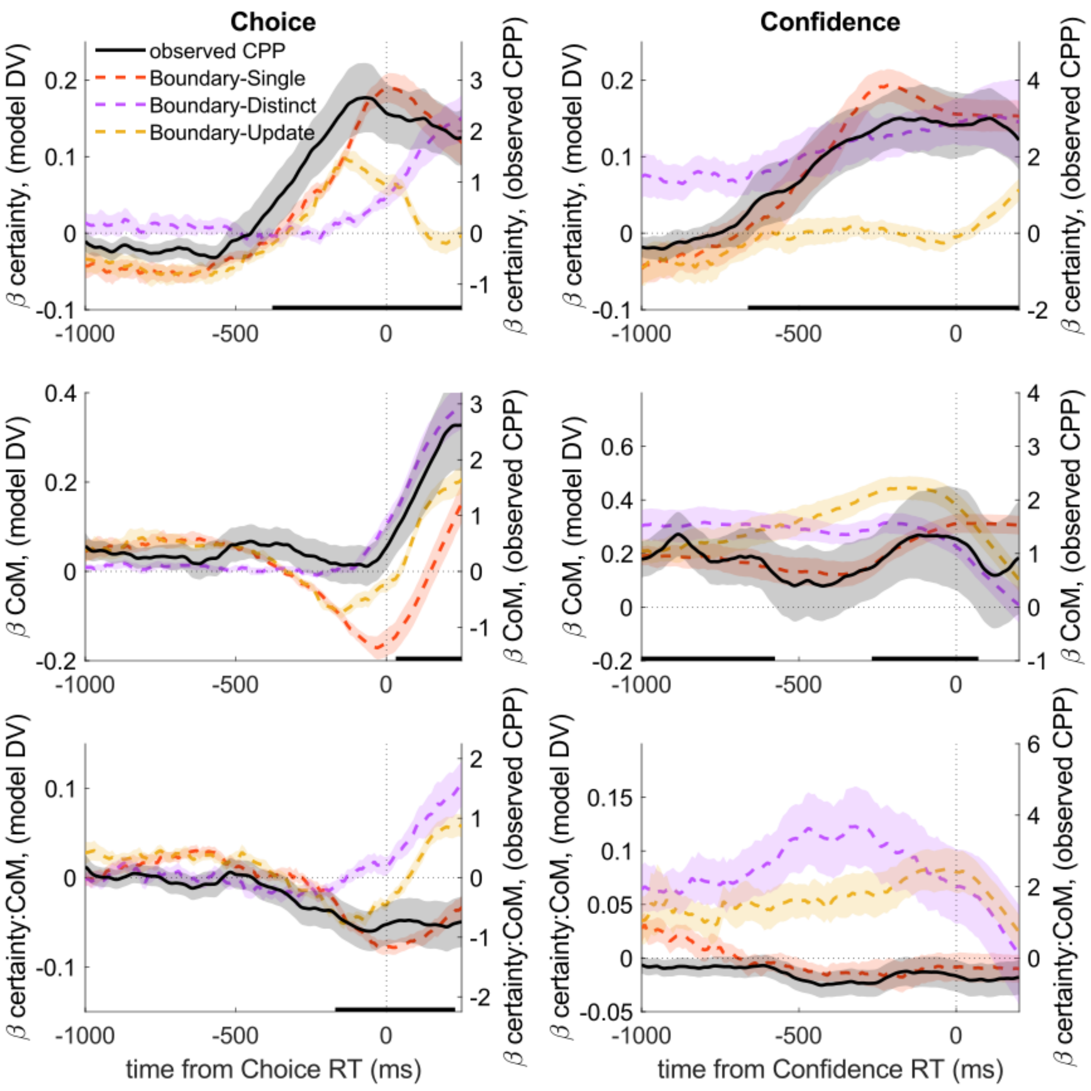


***Extended Data Figure 5. Boundary-Update model variant.*** *This model uses the same parameters as the Boundary-Single model, but here the post-decision process is a separate accumulator that starts from the initial accumulator's end-point (rather than being a continuation of the process). Thus it still carries the same information as the initial-process, but the initial accumulator ends and decays to zero, while the new accumulator begins during the motor execution time. This is similar to the Distinct process, but here the accumulator starts from the initial accumulator's end-point rather than a new starting-point, and the boundaries are set relative to the initial process' starting-points, like in the Single process. This model (Yellow dashed lined) does not capture the Certainty effects, with an initial positive effect decreasing before the initial choice, nor the Certainty*CoM interaction, although the CoM effect does look similar to the CPP.*

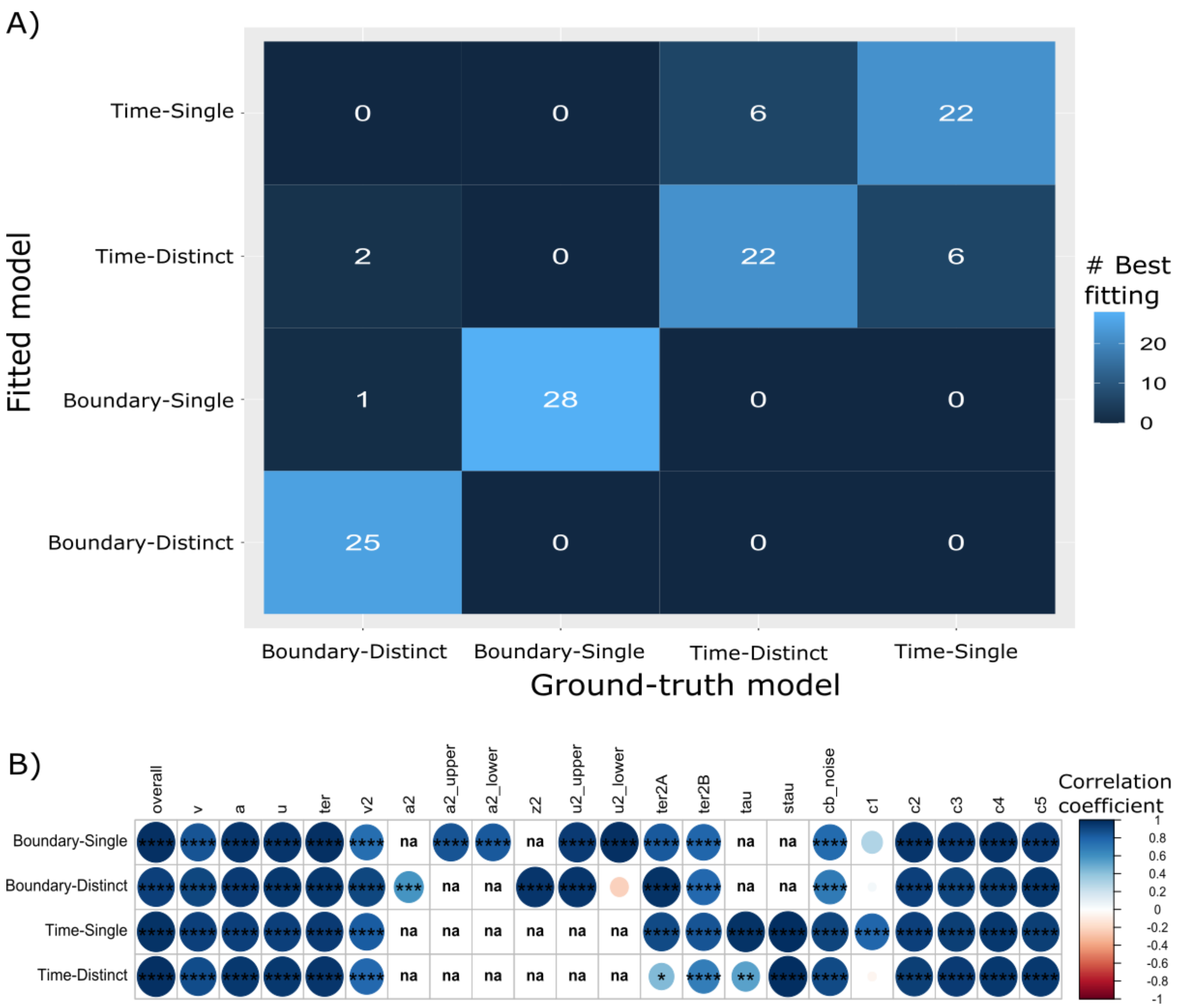


***Extended Data Figure 6. Model and Parameter Recovery plots.*** *A) Model recovery confusion matrix, showing how often the 'ground-truth' model was selected as the best (lowest BIC) when fitting models to data simulated using the parameters fit to behavioural data (per participant & condition). The 'ground-truth' model is recovered most of the time, especially for the Boundary models. B) Parameter recovery for the 'ground-truth' model fits, showing the correlations between the simulated and recovered parameters. 'Overall' refers to correlating all recovered and simulated parameters at once per model, and the other columns refer to separate parameters, with 'na' for parameters not included in a particular model. The size and colour of the circles show the Pearson's correlation coefficient, and the stars show the level of significance (*=$p<.05$, **=$p<.01$, ***=$p<.001$, ***=$p<.0001$). Almost all recovered parameters are strongly correlated, except for 'c1' (1st confidence-criteria) in 3 of the 4 models, and 'u2_lower' for the Boundary-Distinct model.*
